# Multiscale regulation of experience-dependent plasticity by a Pannexin1 homolog in a developing vertebrate brain

**DOI:** 10.64898/2026.04.22.720230

**Authors:** Fatema Nakhuda, Georg S.O. Zoidl, Armin Bahl, Georg R. Zoidl

## Abstract

Experience-dependent plasticity enables the developing brain to adapt to repeated sensory input while maintaining stability. However, how such plasticity is coordinated across behavior, transcription, and network dynamics remains unclear. Here, using larval zebrafish, we identify the pannexin channel homolog, Panx1a as a regulator of multiscale adaptations during visual habituation. Panx1a loss selectively impairs long-term habituation without affecting baseline sensorimotor responses. This behavioral deficit is accompanied by disrupted activity-dependent transcription across distributed brain regions and altered excitation-inhibition balance. At the network level, Panx1a deficiency attenuates experience-dependent modulation of gamma activity, cross-frequency coupling, and inter-regional coherence. We further show that sharp wave-ripple-like events are present in vivo at an early developmental stage and exhibit selective refinement of their sharp-wave component following experience, while ripple features remain largely unchanged. This refinement is reduced in panx1a mutants, indicating a role for Panx1a in shaping network dynamics rather than event generation. Together, these findings identify Panx1a as a mediator linking extracellular signaling to coordinated behavioral, molecular, and circuit-level plasticity during early brain development.

## Introduction

Habituation, the progressive attenuation of a behavioral response to repeated, inconsequential stimuli, is among the most phylogenetically conserved forms of learning (Rankin et al., 2009; Thompson & Spencer, 1966). Far from a passive process of neural fatigue, habituation requires active synaptic modification, including postsynaptic receptor engagement, and at longer timescales, *de novo* gene expression and protein synthesis (Bailey et al., 2015; Cooke et al., 2015; Esdin et al., 2010; Ezzeddine & Glanzman, 2003; Ma et al., 2023). Its defining properties, stimulus specificity, spontaneous recovery, and reversibility by dishabituation, make it a powerful model for studying experience-dependent plasticity (Rankin et al., 2009). Recent work further suggests that habituation reflects an optimization principle, balancing information gain with the metabolic cost of continued responding, implying that neural systems must integrate sensory input with internal energy constraints (Adibi et al., 2013; Attwell & Laughlin, 2001; Nicoletti et al., 2025).

Larval zebrafish provide a tractable vertebrate model for investigating these mechanisms. At 6 days post-fertilization, larvae exhibit robust visually evoked locomotor responses mediated by reticulospinal circuits downstream of the optic tectum (Gahtan et al., 2005; Burgess & Granato, 2007; Randlett et al., 2019; Lamiré et al., 2023). Repeated light stimulation produces progressive response attenuation, and long-term habituation requires NMDA receptor activation and de novo transcription (Roberts et al., 2011, 2016), findings originally established in acoustic and tactile paradigms that parallel molecular requirements identified across other vertebrate systems (Malkani & Rosen, 2000; Mayford et al., 1996; Pellicano et al., 1993; Shimizu et al., 2000). In larval zebrafish, dark-flash habituation, characterized by sudden reductions in illumination, persists for multiple hours and depends on GABAergic inhibition, with distributed plasticity engaging tectal and pallial circuits (Lamiré et al., 2023; Randlett et al., 2019), Despite these advantages, the upstream signaling pathways linking neural activity to transcriptional responses during habituation remain poorly understood.

Pannexin 1 (Panx1) is a membrane channel widely expressed in the vertebrate brain (Ray et al., 2005; Vogt et al., 2005; G. Zoidl et al., 2007) and enriched at postsynaptic sites of excitatory synapses (G. Zoidl et al., 2007). Its primary function is the release of ATP (Bao et al., 2004; Dahl, 2015), which initiates purinergic signaling through P2X, P2Y, and adenosine (P1) receptors (Khakh & North, 2012; Latini & Pedata, 2001; North & Verkhratsky, 2006). This signaling cascade modulates neurotransmission, intracellular calcium, and gene expression across neurons and glia (Mayhew et al., 2018; Pankratov et al., 2009; Shigetomi et al., 2024). At the synaptic level, Panx1 interacts with NMDA receptor signaling and in turn modulates NMDAR-dependent currents through purinergic feedback (Patil et al., 2022; Rangel-Sandoval et al., 2024a; Weilinger et al., 2016). Loss of Panx1 alters GABAergic transmission, modifies synaptic plasticity thresholds (Ardiles et al., 2014; Gajardo et al., 2018; García-Rojas et al., 2023), and affects dendritic structure (Flores-Muñoz et al., 2022; Sanchez-Arias et al., 2019, 2020) and network synchronization (Sanchez-Arias et al., 2019, p. 20). Consistent with these roles, Panx1 deficiency has been linked to impairments in learning and memory across model systems (Casillas Martinez et al., n.d.; Gajardo et al., 2018; Illanes-González et al., 2025; Obot et al., 2023, 2024; Prochnow et al., 2012). However, how Panx1 influences brain-wide transcriptional programs and network-level dynamics during learning remains unknown.

Neural oscillations provide a systems-level framework for understanding coordinated circuit activity during learning. Theta and gamma oscillations, and their coupling, are closely associated with memory processes (Kendrick et al., 2011; Tort et al., 2009; Wulff et al., 2009), while gamma activity reflects interactions between excitatory neurons and inhibitory interneurons (Amilhon et al., 2015; Buzsáki & Wang, 2012; Fries, 2015). Disruptions in these dynamics often indicate altered excitation-inhibition balance. Sharp wave-ripple (SWR) complexes, characterized by large-amplitude slow waves accompanied by transient high-frequency oscillations, are a hallmark of offline hippocampal activity associated with memory consolidation (Buzsáki, 2015; Colgin, 2016; Koniaris et al., 2011). In zebrafish, recent work has identified sharp wave-ripple-like events in hippocampal homologs, suggesting conserved mechanisms of network coordination (Blanco et al., 2024). Whether such dynamics are present during early development in vivo, and how they are modulated by experience, remains unclear.

Here, we use a *panx1a* loss-of-function zebrafish model (*panx1a^−/−^*, TALEN-generated premature stop) to examine the role of Pannexin1 channel in visual habituation. We show that *panx1a* is broadly expressed across the larval brain and that its loss produces selective neuroanatomical alterations. Behaviorally, *panx1a^−/−^* larvae exhibit impaired long-term habituation while maintaining baseline sensorimotor responses, indicating a specific deficit in experience-dependent plasticity. This impairment is accompanied by disrupted brain-wide transcriptional responses and altered excitation-inhibition balance, as revealed by pharmacological manipulations. At the network level, Panx1a loss reduces gamma power, disrupts theta-gamma coupling, and alters inter-regional coordination. We further demonstrate that sharp wave-ripple-like events are present in vivo at larval stages, are modulated by experience, and depend on Panx1a. Together, these findings identify Panx1a as a multiscale regulator linking extracellular purinergic signaling to transcriptional and network-level plasticity during learning.

## Results

### Panx1a is broadly expressed in the larval zebrafish brain and is required for visual habituation

We first characterized the spatial distribution of *panx1a* transcript in 6-dpf WT zebrafish larvae using Hybridization Chain Reaction RNA fluorescence in situ hybridization (HCR RNA-FISH). Panx1a mRNA was widely expressed throughout the brain, with prominent signal in the optic tectum (TeO), telencephalon, and hindbrain **(Fig. 1A–C)**. Quantification of fluorescence intensity across annotated brain regions revealed region-specific differences in expression, including a rostro–caudal gradient along the anterior–posterior axis **(Fig. 1B, C)**. Panx1a signal was observed in proximity to postsynaptic density (PSD) markers, indicating localization within synaptic regions **(Fig. 1D, top panel)**. In addition, *panx1a* transcript expression was distributed across regions corresponding to both glutamatergic (slc-defined) and GABAergic (gad-defined) domains in the MapZebrain atlas, suggesting broad positioning across excitatory and inhibitory systems **(Fig 1D, bottom panel)**.

**Figure 1.**
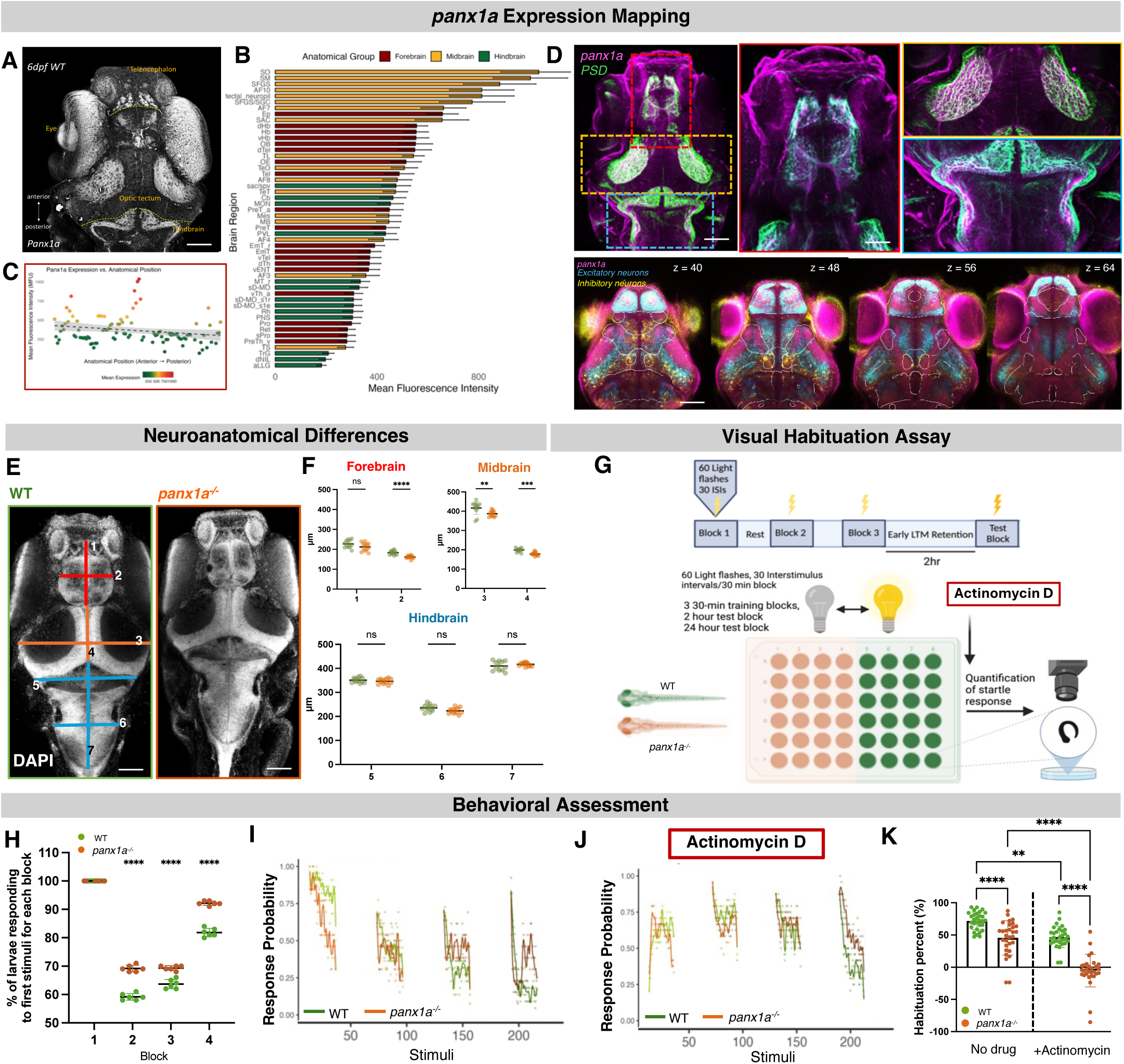
*panx1a* expression, neuroanatomical alterations, and impaired visual habituation in larval zebrafish. **(A–C)** Whole-brain mapping of *panx1a* mRNA expression in 6-dpf WT larvae using HCR RNA-FISH. **(A)** Dorsal view showing widespread expression across major brain regions, including telencephalon, optic tectum (TeO), and hindbrain. **(B)** Quantification of mean fluorescence intensity across annotated brain regions, grouped by major anatomical divisions, revealing region-specific enrichment. **(C)** Distribution of *panx1a* expression along the anterior–posterior axis, showing a rostro–caudal gradient. **(D)** High-resolution confocal images showing *panx1a* signal (magenta) with neuronal marker co-labeling (green), indicating expression within neuronal populations. Insets highlight regional localization across z-planes. **(E–F)** Neuroanatomical analysis in WT and *panx1a^−/−^* larvae. **(E)** Representative DAPI-stained brains with standardized measurement axes. **(F)** Quantification of regional brain dimensions reveals reduced forebrain width and optic tectum size in *panx1a^−/−^* larvae, with no changes in hindbrain measurements. **(G)** Schematic of the visual habituation paradigm consisting of repeated light stimuli across training blocks followed by retention testing. **(H–I)** Habituation dynamics across training blocks. Percentage of responding larvae to initial and repeated stimuli (stimulus number shown on the y-axis) shows progressive response reduction in WT and impaired habituation in *panx1a^−/−^* larvae. **(J)** Effect of transcriptional inhibition (Actinomycin D) on habituation dynamics. **(K)** Quantification of habituation percentage in WT and *panx1a^−/−^* after adding ActinomycinD. Data are presented as mean ± SD. Statistical significance was determined using two-way ANOVA with Šídák’s multiple comparisons test and is indicated (*p < 0.05, **p < 0.01, ****p < 0.0001; ns, not significant).

To assess whether loss of Panx1a loss alters brain size, we performed DAPI-based morphometric analysis in WT and *panx1a^−/−^* larvae at 6-dpf. Panx1a-deficient larvae exhibited selective reductions in forebrain width (WT: 183.2 µm; *panx1a^−/−^* : 161.1 µm; p < 0.0001) and optic tectum dimensions, including decreased length (WT: 416.4 µm; *panx1a^−/−^*: 386.8 µm; p = 0.0023) and width (WT: 198.5 µm; *panx1a^−/−^*: 178.5 µm; p = 0.0097) **(Fig. 1E, F; Supplementary Table 1)**. In contrast, hindbrain measurements were unchanged, indicating that Panx1a loss results in region-specific anatomical alterations rather than global developmental defects.

We next examined the functional consequences of Panx1a loss using a light-flash habituation paradigm **(Fig. 1G)**, in which repeated visual stimuli elicit progressively reduced behavioral responses. WT larvae exhibited robust within-session habituation across repeated stimuli, with response probability progressively decreasing over successive stimuli and across training blocks, and remaining suppressed at the 2 h test time point **(Fig. 1H–K)**. At the block level, WT larvae showed a marked reduction in the proportion of responders after the first stimulus across successive blocks, consistent with effective habituation **(Fig. 1H)**.

In contrast, *panx1a^−/−^* larvae showed impaired habituation, with weaker response attenuation during training and elevated response probability at 2 h **(Fig. 1H–K; Supplementary Table 1)**. This impairment was evident across blocks, where mutants maintained a higher proportion of responders following repeated stimulation **(Fig. 1H)**. Notably, baseline responses to the initial stimulus were comparable between genotypes, indicating intact sensory detection and motor output.

To determine whether transcription is required for habituation, we inhibited transcription using Actinomycin D. In WT larvae, transcriptional blockade significantly reduced habituation **(Fig. 1J, K)**. In *panx1a^−/−^* larvae, Actinomycin D further disrupted habituation relative to their already impaired baseline **(Fig. 1J, K; Supplementary Table 1)**. Given that WT larvae exhibit stable retention at 2 h while mutants already display pronounced deficits, subsequent analyses focused on this time point to probe the molecular and circuit mechanisms underlying Panx1a-dependent plasticity.

### Habituation engages distributed transcriptional programs that require Panx1a

To determine how habituation modulates transcription, we first assessed activity-dependent gene expression using targeted analysis of IEGs. In WT larvae, habituation induced robust expression of IEGs, including *fosab*, consistent with activation of transcriptional programs following sensory experience **(Fig. 2B)**. In contrast, *panx1a^−/−^* larvae showed reduced or altered expression across multiple IEGs, indicating impaired activity-dependent transcriptional responses.

**Figure 2.**
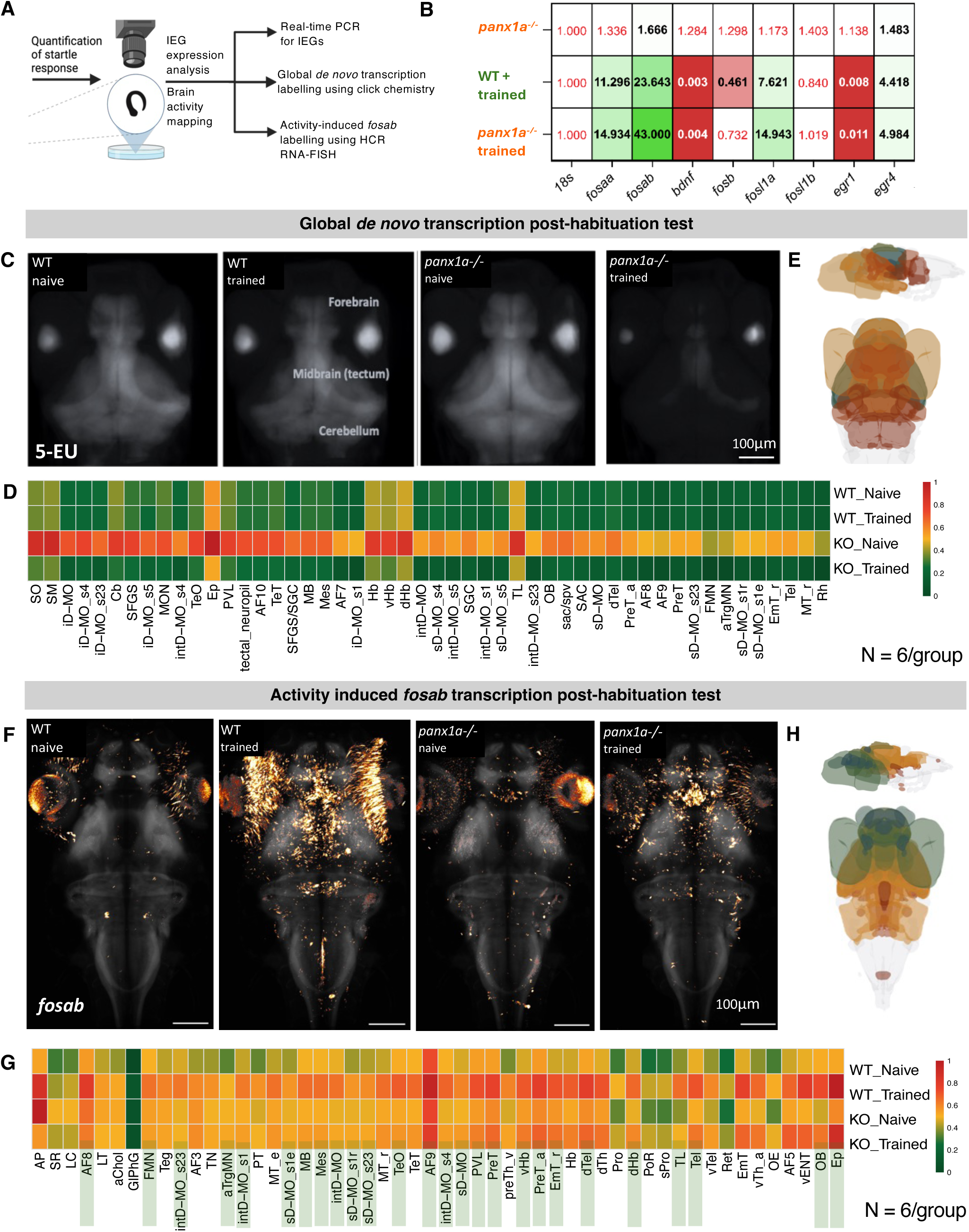
Panx1a-dependent transcriptional responses to visual habituation. **(A)** Experimental workflow linking behavioral habituation to transcriptional and activity-dependent readouts, including de novo RNA synthesis (5-EU labeling) and *fosab* expression. Values in red text are non-significant. **(B)** Heatmap of IEG expression across knockout and trained conditions, highlighting differential regulation following habituation. All fold change values are normalized to naïve WT. **(C)** Representative whole-brain 5-EU labeling in WT and *panx1a^−/−^* larvae under naïve and trained conditions. Scale bar, 100μm **(D)** Quantification of regional 5-EU signal intensity across anatomically defined brain regions. **(E)** 3D mapping of transcriptional changes across brain regions, color-coded by training-induced effect size. **(F)** Whole-brain HCR RNA-FISH detection of *fosab* expression in WT and *panx1a^−/−^* larvae under naïve and trained conditions. Scale bar, 100μm. **(G)** Quantification of *fosab* signal intensity across brain regions, showing reduced activity-dependent induction in *panx1a^−/−^* larvae. **(H)** 3D reconstruction of *fosab*-positive signal across brain regions, color-coded by effect strength. Data are presented as mean values (N = 6 larva per group). Statistical details are provided in Supplementary Tables 3-4.

We next quantified *de novo* RNA synthesis using 5-ethynyl uridine (5-EU) incorporation across anatomically defined regions. We focused our analysis on regions that showed robust signal and consistent changes across both transcriptional and activity-dependent readouts **(Supplementary Tables 2–3).**

In WT larvae, habituation was associated with coordinated transcriptional activity across multiple brain regions, including the optic tectum (TeO), cerebellum (Cb), habenula (dHb, vHb), and epithalamus (Ep) **(Fig. 2C–E)**. These regions are associated with sensory processing, integration, and behavioral adaptation. Across these areas, transcriptional responses were largely maintained following training, suggesting stable engagement of distributed brain networks.

In contrast, *panx1a^−/−^* larvae showed elevated baseline transcription across many of these same regions that was not sustained after habituation. Instead, transcriptional signal decreased following training, notably in optic tectum, cerebellum, epiphysis, dorsal habenula **(Fig. 2C–E; Supplementary Table 3).** Similar patterns were observed in hindbrain sensorimotor nuclei and tectal neuropil layers, pointing to a broad disruption of activity-dependent transcription.

To assess activity-dependent gene expression, we quantified *fosab* using HCR RNA-FISH. WT larvae showed robust induction of *fosab* following habituation across forebrain, midbrain, and hindbrain regions, including the optic tectum, habenula, dorsal telencephalon, and epiphysis **(Fig. 2F–H; Supplementary Table 4)**. In contrast, *panx1a^−/−^* larvae showed weaker or absent induction, and in some regions, reduced signal after training (e.g., area postrema: 1624.0 → 1000.4; superior raphe: 906.1 → 806.1). Regions that showed strong induction in WT, including epiphysis and dorsal telencephalon, displayed only partial responses in mutants.

Across regions, the patterns observed with 5-EU and *fosab* were consistent, indicating a failure to sustain activity-transcription coupling in the absence of a functional Panx1a. Together, these results show that habituation engages coordinated, brain-wide transcriptional responses, and that Panx1a is required to maintain these responses across functionally relevant regions.

### Panx1a regulates visual habituation through modulation of excitatory and inhibitory signaling

To test whether Panx1a influences circuit excitability during habituation, we pharmacologically manipulated glutamatergic and GABAergic signaling during the visual habituation assay **(Fig. 3A).** Under control conditions, *panx1a^−/−^* larvae exhibited impaired habituation relative to WT, as indicated by reduced attenuation of responses across repeated stimuli **(Fig. 3B–F; Supplementary Table 1).**

**Figure 3.**
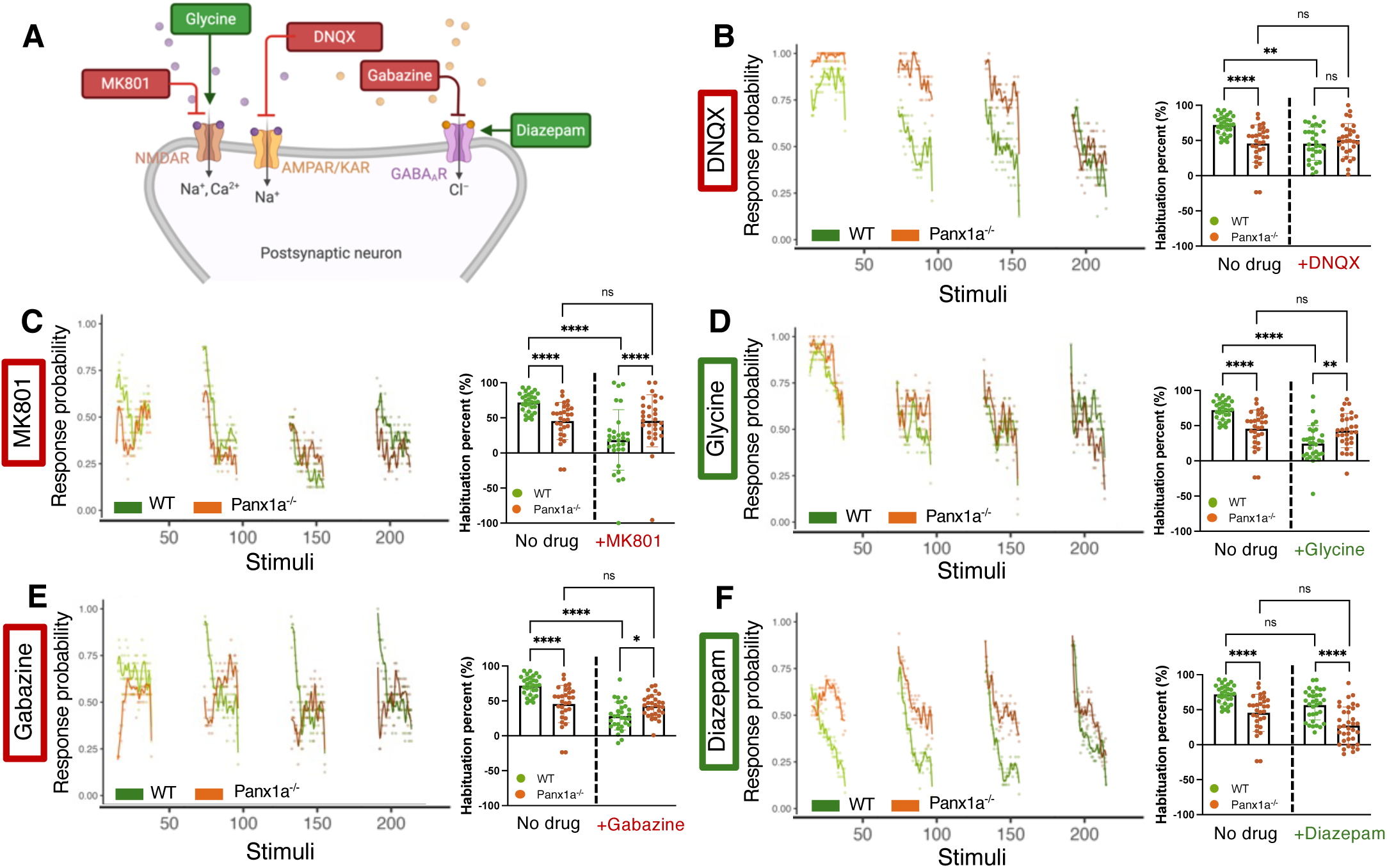
Panx1a-dependent modulation of excitation–inhibition balance during visual habituation. **(A)** Schematic of pharmacological manipulations during the visual habituation assay targeting glutamatergic and GABAergic signaling pathways. **(B)** Effects of AMPA receptor blockade (DNQX) on habituation. Left: trial-by-trial response probability across repeated stimuli. Right: quantification of habituation percentage under control and DNQX conditions. **(C)** Effects of NMDA receptor antagonism (MK-801) on habituation dynamics and overall response attenuation. **(D)** Effects of NMDA receptor potentiation with glycine on habituation across genotypes. **(E)** Effects of GABA_A_ receptor inhibition (gabazine) on habituation. **(F)** Effects of GABA_A_ receptor potentiation (diazepam), showing no rescue of habituation deficits in *panx1a^−/−^* larvae. Across panels, line plots represent mean response probability across repeated stimuli, and bar graphs summarize habituation percentage. Data are presented as mean ± SD. Statistical significance was determined using two-way ANOVA with Šídák’s multiple comparisons test and is indicated (*p < 0.05, **p < 0.01, ****p < 0.0001; ns, not significant).

We systematically probed excitatory and inhibitory signaling to determine whether Panx1a-dependent habituation deficits arise from altered synaptic transmission, plasticity, or inhibitory control. We first examined glutamatergic signaling, targeting AMPA receptors to assess fast excitatory transmission and NMDA receptors to probe activity-dependent plasticity. We then manipulated GABAergic signaling to test whether inhibitory control of neural activity contributes to habituation and the observed mutant phenotype. Blockade of AMPA receptors with DNQX reduced overall habituation in WT larvae and diminished genotype-dependent differences **(Fig. 3B)**, indicating that fast glutamatergic transmission contributes to habituation dynamics. Similarly, NMDA receptor antagonism with MK-801 markedly reduced habituation in WT larvae while producing minimal additional effects in *panx1a^−/−^* mutants, thereby reducing genotype-dependent differences **(Fig. 3C).** These results are consistent with a role for NMDAR-dependent plasticity in habituation.

To further probe NMDA receptor involvement, we enhanced NMDAR function using glycine. Glycine reduced habituation in WT larvae but had limited effects in *panx1a^−/−^* mutants, revealing differential sensitivity to NMDAR modulation **(Fig. 3D).**

We next examined inhibitory signaling. Blockade of GABA_A_ receptors with gabazine disrupted habituation in both genotypes and attenuated genotype-dependent differences **(Fig. 3E),** indicating that inhibitory transmission is required for normal habituation. In contrast, potentiation of GABA_A_ receptors with diazepam did not significantly change habituation in either genotype and did not rescue the deficit observed in *panx1a^−/−^* larvae **(Fig. 3F)** suggesting that increasing inhibitory tone alone is insufficient to restore Panx1a-dependent learning deficits. Across manipulations, excitatory and inhibitory perturbations produced complementary effects on habituation. Notably, increasing inhibitory tone alone was insufficient to restore the mutant phenotype, indicating that the observed deficits do not arise from a simple reduction in inhibition but reflect a broader imbalance in circuit dynamics. Similar effects were observed across additional pharmacological manipulations targeting the same pathways **(Supplementary Fig. 2)**, indicating that these results are robust across independent compounds. Together, these findings indicate that Panx1a regulates habituation by modulating circuit-level excitation–inhibition balance.

### Panx1a regulates experience-dependent oscillatory dynamics and network coordination during habituation

Given the behavioral and transcriptional abnormalities observed in *panx1a^−/−^* larvae, we next examined whether network-level activity is altered during habituation. Local field potentials (LFPs) were recorded simultaneously from the antero-dorsolateral pallium (ADL), a higher-order associative region, and the optic tectum (TeO), a primary visual processing center, in 6-dpf WT and *panx1a^−/−^* larvae in vivo **(Fig. 4A)**. Representative traces illustrate theta (3-7 Hz) and gamma (30-45 Hz) activity across conditions **(Fig. 4B)**.

**Figure 4.**
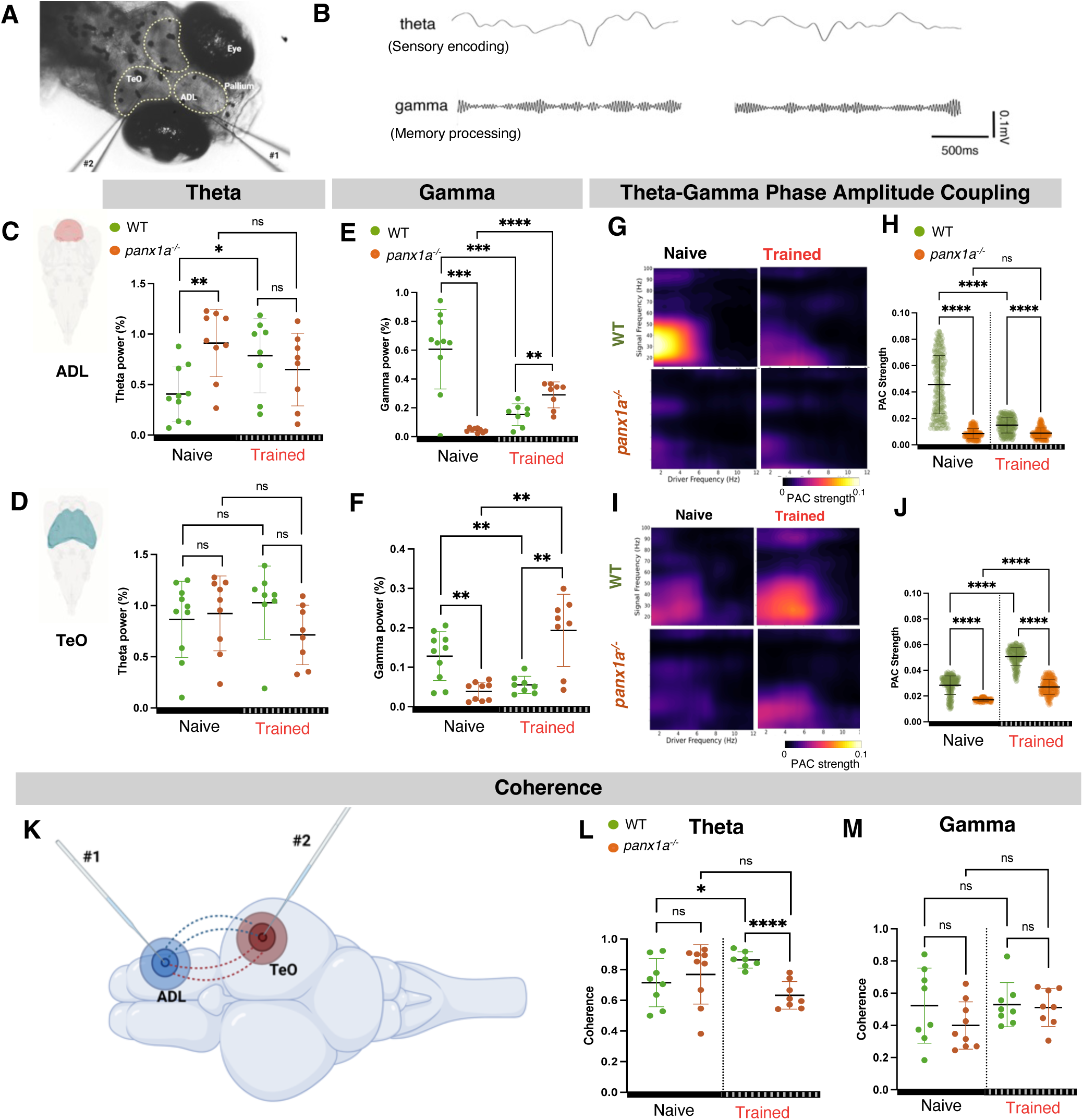
Panx1a regulates oscillatory dynamics and cross-frequency coupling during visual habituation. **(A–B)** Electrophysiology overview. **(A)** Schematic of the electrophysiology setup indicating dual electrode placement for recording brain activity in vivo. **(B)** Representative traces showing theta-band activity (sensory encoding) and gamma-band activity (memory processing). **(C–F)** Region-specific oscillatory power changes. **(C–D)** Theta power in the ADL and TeO in WT and *panx1a^−/−^* larvae under naïve and trained conditions. Training modestly increased the theta-band power of WT larvae. **(E–F)** Gamma power in ADL and TeO. WT larvae show decreased gamma power following habituation, whereas *panx1a^−/−^* larvae exhibit increased gamma responses across both brain regions. **(G–J)** Theta-gamma PAC. **(G, I)** Comodulograms illustrating PAC strength across frequency pairs in ADL and TeO for naïve and trained conditions. WT larvae show enhanced PAC following training, especially in the TeO. **(H, J)** Quantification of theta-gamma PAC strength confirms training-induced decreases in the ADL of WT larvae, while significant training-induced increases in WT that are diminished in *panx1a^−/−^* larvae are observed in the TeO. **(K–M)** Coherence analysis. **(K)** Schematic illustrating dual-site LFP recording schematic illustrating inter-regional coherence between ADL and TeO. Electrodes (#1, #2) capture local activity (concentric rings), while synchronized oscillations (dashed lines) indicate functional coupling across regions. **(L–M)** Coherence measurements between the ADL and TeO show frequency-specific alterations following habituation. WT larvae exhibit increased theta-band coherence after training, whereas *panx1a^−/−^* larvae show reduced or disrupted coherence compared to the WT; no significant changes are observed in the gamma-band. Data are presented as individual data points with mean ± SD. Number of animals: WT, WT Trained, panx1a^−/−^ trained N = 8; *panx1a^−/−^* N = 9. Outliers were removed using the ROUT test (Q = 1%). Data were analyzed using Welch’s t-test (PSD and coherence) and a two-way ANOVA with Tukey’s multiple comparisons test (PAC). Statistical significance is indicated as *p < 0.05, **p < 0.01, ***p<0.001, ****p < 0.0001, ns, not significant.

Theta-band power did not show consistent training-dependent changes across genotypes or regions **(Fig. 4C, D)**, indicating that low-frequency activity is largely preserved. In contrast, gamma-band activity was selectively modulated by experience. WT larvae exhibited training-dependent reductions in gamma power, particularly in the ADL, whereas *panx1a^−/−^* larvae showed blunted modulation in both ADL and TeO **(Fig. 4E, F)**, indicating impaired regulation of higher-frequency oscillations.

We next assessed theta-gamma phase-amplitude coupling (PAC), a measure of cross-frequency coordination. In WT larvae, habituation decreased PAC in the ADL, as reflected in both comodulograms and quantification **(Fig. 4G, H)**. This decrease was absent in *panx1a^−/−^* larvae. In the TeO, PAC was significantly increased by training across genotypes **(Fig. 4I, J)**, indicating region-specific effects.

Finally, we examined inter-regional coherence as the TeO and ADL are part of the ascending visual pathway **(Fig. 4K)**. WT larvae showed increased coherence in the theta-band following habituation, whereas *panx1a^−/−^* larvae exhibited reduced or disrupted coherence compared with trained WT **(Fig. 4L)**. In contrast, coherence in the gamma-band remained largely unaffected by training across conditions **(Fig. 4M)**, suggesting that Panx1a preferentially supports network coordination in slower-wave coherence. Collectively, these results show that Panx1a is required for experience-dependent modulation of gamma activity, cross-frequency coupling, and network coherence. Rather than reflecting a global loss of oscillatory activity, Panx1a deficiency selectively disrupts higher-frequency and integrative network dynamics that support learning.

### Experience-dependent modulation of sharp wave-ripple dynamics in the zebrafish pallium requires Panx1a

To determine whether experience-dependent plasticity extends to transient network events, we analyzed sharp wave-ripple (SWR)-like complexes recorded from the dorsolateral pallium (ADL) of 6-dpf larvae in vivo **(Fig. 5A, B)**. These events consisted of a large-amplitude sharp-wave (SW) deflection temporally coupled to a brief burst of high-frequency ripple activity, consistent with canonical SWR signatures. Time-frequency analysis aligned to ripple peaks confirmed a transient increase in ripple-band power, supporting the identification of these events as SWR-like complexes.

**Figure 5.**
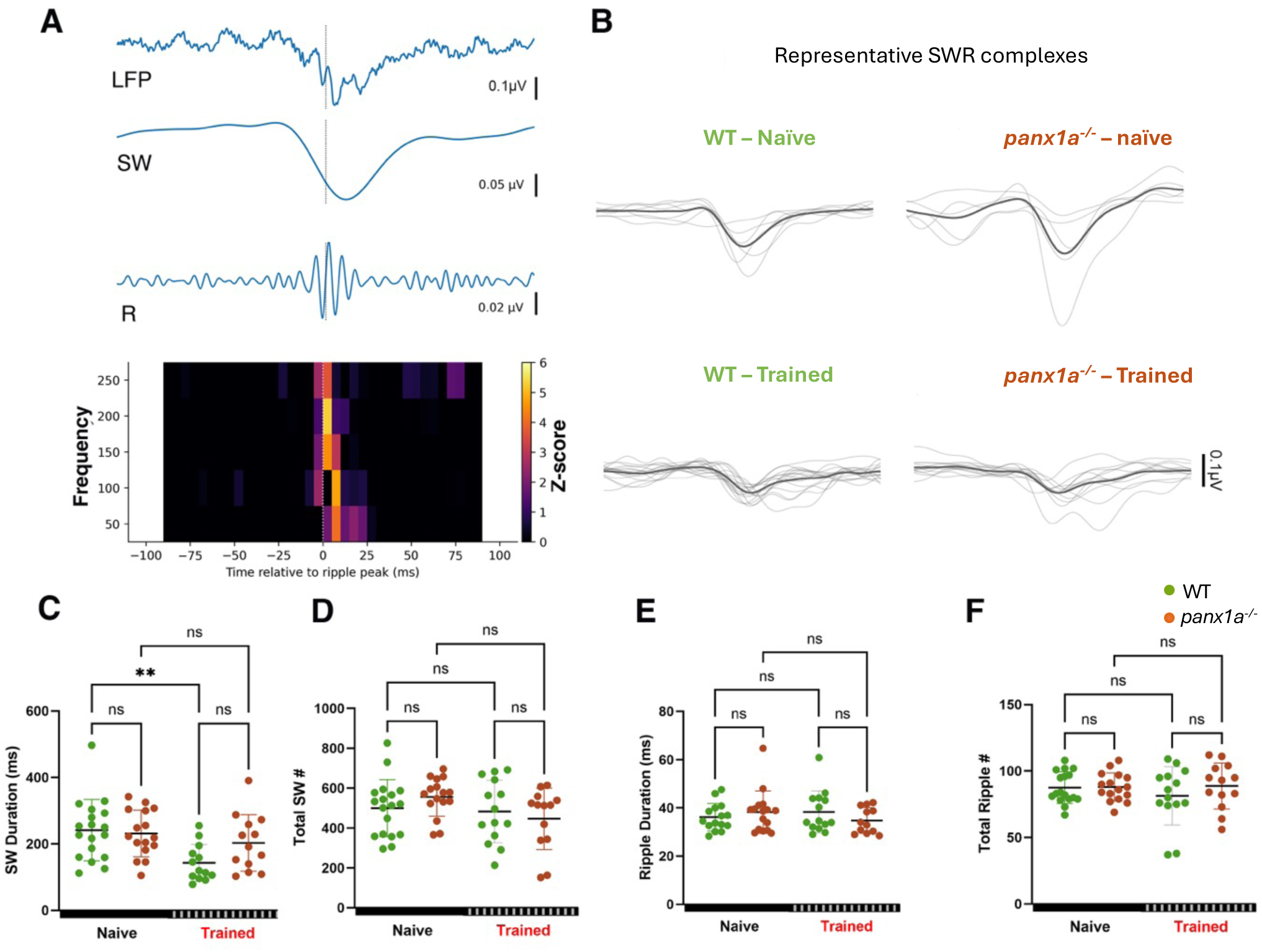
Panx1a regulates experience-dependent SWR duration in the dorsolateral pallium. **(A)** Representative SWR-like complex recorded in vivo from the ADL of 6-dpf larvae. The LFP shows a large-amplitude SW coincident with a transient high-frequency ripple (R) burst. Time-frequency spectrogram aligned to ripple peak (dashed line) confirms a temporally restricted increase in ripple-band power. **(B)** Representative SW waveforms for WT and *panx1a^−/−^* larvae under naïve and trained conditions (gray, individual events; black, mean), illustrating experience-dependent shortening in WT and increased variability in mutants. **(C–F)** Quantification of SWR features (points represent individual larvae; mean ± SD). **(C)** SW duration is reduced following training in WT larvae; an effect attenuated in *panx1a^−/−^*. **(D)** Total number of SW events shows no significant training or genotype effect under mixed-model analysis. **(E)** Ripple duration is not significantly altered across conditions. **(F)** Total ripple count remains unchanged across genotype and training. Outliers were removed using the ROUT test (Q = 1%). Statistical comparisons were performed using a two-factor linear model testing the effects of genotype, treatment, and their interaction, with Tukey-adjusted post hoc comparisons of estimated marginal means. Number of animals: WT naïve N = 18; WT Trained N = 14; *panx1a^−/−^* naïve N = 16; *panx1a^−/−^*trained N = 13. Significance is indicated as **P < 0.01, ns, not significant.

Visual habituation was associated with a selective modification of SW structure in the ADL of WT larvae. Specifically, the SW duration was reduced following training, indicating experience-dependent refinement of sharp-wave temporal dynamics **(Fig. 5C)**. In contrast, this shortening was attenuated in *panx1a^−/−^* larvae, in which SW waveforms exhibited increased variability and lacked consistent training-induced modulation. These observations were confirmed using a two-factor linear model, which revealed a significant effect of training on SW duration in WT animals (P<0.01), but not in Panx1a^−/−^ mutants, resulting in a genotype-dependent divergence after training.

In contrast to these changes in sharp-wave structure, other SWR features were largely preserved **(Fig. 5D-F)**. The total number of SW events did not differ significantly across genotype or training conditions, nor did ripple duration or total ripple count. These results indicate that the generation and frequency of SWR-like events remain intact in the absence of Panx1a, whereas the temporal structure of the sharp-wave component is selectively sensitive to experience.

Together, these findings show that SWR-like activity is present in the developing zebrafish pallium and is modulated by sensory experience. Panx1a is not required for the occurrence of these events per se but is necessary for their experience-dependent temporal refinement, suggesting a role in shaping network-level dynamics associated with early forms of learning.

## Discussion

Pannexin 1 channels have been implicated in synaptic signaling, circuit development, and purinergic communication in the central nervous system. Through ATP release and downstream receptor activation, Panx1 is positioned to influence both neuronal excitability and activity-dependent plasticity. However, how these functions contribute to learning-related processes across behavioral, molecular, and network levels remains incompletely understood. Here, we identify Panx1a as a regulator of experience-dependent plasticity in larval zebrafish, linking habituation behavior to coordinated transcriptional and circuit-level dynamics.

### Panx1a loss selectively disrupts experience-dependent plasticity

Panx1a is broadly expressed across the larval zebrafish brain, with enrichment in sensory and integrative regions such as the optic tectum (TeO) and telencephalon. In mammals, PANX1 has been shown to regulate neural progenitor proliferation and differentiation through ATP-mediated purinergic signaling, suggesting that its loss can produce region-specific developmental and circuit-level effects rather than global disruption (Wicki-Stordeur & Swayne, 2013). Consistent with this, *panx1a^−/−^* larvae exhibit selective reductions in forebrain and tectal dimensions, pointing to localized vulnerability in circuits that process and integrate sensory information.

The reduction in tectal structure is notable given that our HCR RNA-FISH data place *panx1a* transcript near a synaptic marker. Previous work has localized Panx1 protein to postsynaptic compartments and demonstrated its association with PSD-95 in mammalian neurons (G. Zoidl et al., 2007), supporting a role in synaptic signaling. In this context, Panx1a-mediated ATP release may contribute to the stabilization or modulation of active synapses, particularly in circuits that undergo repeated sensory engagement. Its absence may therefore compromise the structural or functional integrity of high-demand networks, contributing to the regional anatomical differences observed in mutants.

At the behavioral level, *panx1a^−/−^* larvae display a selective impairment in habituation despite preserved baseline locomotor capacity (Figure. 1I, Supp. Figure. S1). In a no-stimulus condition, mutants exhibit activity levels comparable to WT, indicating that the observed phenotype is unlikely to arise from gross motor deficits. While mutants show some response decrement during initial stimulation, they fail to maintain suppression at the 2-hour test point. This pattern is consistent with a disruption in the stabilization or consolidation of habituation rather than in sensory detection or motor output.

Interestingly, independent observations from our lab indicate that *panx1a^−/−^* larvae exhibit reduced and intermittent locomotor activity during the dark phase, characterized by bouts of slow- and medium-speed swimming (Safarian et al., 2020). Together, these findings suggest that Panx1a does not regulate baseline locomotion per se but may instead contribute to the context-dependent modulation of behavioral state. In this framework, the habituation deficit reflects a failure to appropriately adjust behavior in response to repeated sensory input, rather than an inability to generate or sustain movement. In zebrafish and other systems, long-term habituation depends on spaced training and requires new gene expression and protein synthesis (Esdin et al., 2010; Ezzeddine & Glanzman, 2003; Rankin et al., 2009; Roberts et al., 2016). Supporting this, transcriptional blockade in WT larvae phenocopies aspects of the *panx1a^−/−^* deficit, suggesting that Panx1a contributes to processes that sustain experience-dependent behavioral change.

### Purinergic signaling and inhibitory regulation during habituation

One mechanism that may underlie the failure to maintain habituation is impaired purinergic feedback. Panx1 channels are a major pathway for ATP release, and extracellular ATP can be converted to adenosine, which generally suppresses neuronal excitability through adenosine receptor signaling (Burnstock et al., 2010; Dahl, 2015). During repeated stimulation, such signaling may contribute to activity-dependent dampening of circuit output. In the absence of Panx1a, this feedback loop may be weakened, allowing circuits to recover responsiveness more rapidly and preventing stable suppression. This interpretation is consistent with broader principles of homeostatic plasticity, in which neural circuits adjust excitability to maintain stable function (Pozo & Goda, 2010; Turrigiano, 2008). In *panx1a^−/−^* larvae, the failure to sustain response suppression may reflect a disruption in excitation–inhibition balance. Panx1 regulates GABAergic transmission, and its loss reduces inhibitory efficacy while shifting circuits toward excitation (García-Rojas et al., 2023), a change that could impair activity-dependent dampening during repeated stimulation.

### Disruption of activity-dependent transcriptional programs

At the molecular level, Panx1a loss disrupts activity-dependent transcriptional responses, effectively decoupling sensory experience from sustained gene expression changes. In WT larvae, habituation is associated with coordinated increases in de novo RNA synthesis and immediate early gene (IEG) expression across multiple brain regions. In contrast, *panx1a^−/−^* larvae show a blunted and less organized transcriptional response following training.

Activity-dependent transcription in neurons is typically driven by calcium-dependent signaling pathways that link synaptic activity to transcription factor activation and IEG induction (Greer & Greenberg, 2008; Lyons & West, 2011). Panx1-mediated ATP release has been shown to influence intracellular calcium dynamics through purinergic receptor activation (Dahl, 2015; Locovei et al., 2006; Swayne & Boyce, 2017), suggesting a potential mechanism by which Panx1a could contribute to activity-transcription coupling. Through this pathway, Panx1-mediated signaling has the potential to influence calcium-dependent transcriptional responses. In this framework, Panx1a may contribute to activity–transcription coupling by facilitating the translation of synaptic activity into sustained gene expression programs. Interestingly, *panx1a^−/−^* larvae exhibit elevated baseline transcription in several regions. This may reflect compensatory circuit regulation, as neural systems are known to adjust baseline excitability in response to perturbations to maintain functional stability (Pozo & Goda, 2010; Turrigiano, 2008). An elevated baseline state could reduce the dynamic range available for further activity-dependent induction, effectively imposing a ceiling on transcriptional responses. Activity-dependent gene expression is known to depend on stimulus intensity and intracellular signaling thresholds (Greer & Greenberg, 2008; Tyssowski et al., 2018), suggesting that elevated baseline activity may constrain the ability of circuits to mount coordinated transcriptional responses during habituation.

### Panx1a and excitation–inhibition balance

Our pharmacological data further suggest that Panx1a contributes to the regulation of excitation–inhibition (E/I) balance during habituation. NMDA receptor antagonism reduces habituation in WT larvae but produces minimal additional effects in *panx1a^−/−^* mutants, consistent with partial convergence between Panx1a-dependent signaling and NMDAR-linked plasticity mechanisms. Panx1 and NMDAR signaling are functionally coupled in mammalian neurons, with NMDAR-mediated calcium influx triggering Panx1 channel activation and Panx1-dependent ATP release further modulating excitability and synaptic signaling (Li et al., 2018; Rangel-Sandoval et al., 2024b; Weilinger et al., 2016). In this framework, loss of Panx1 disrupts this feedback loop, which may alter NMDAR-dependent activity and impair the regulation of circuit excitability.

Perturbation of GABAergic signaling strongly affects habituation, yet enhancement of GABA_A_ receptor function does not rescue the mutant phenotype. This argues against a simple reduction in inhibitory tone and instead suggests a defect in the activity-dependent recruitment or regulation of inhibition during repeated stimulation. This interpretation is consistent with evidence that visual habituation in larval zebrafish is sensitive to GABA_A_/GABA_C_ receptor blockade, indicating that inhibitory signaling contributes to habituation learning, and with broader work showing that habituation and experience-dependent sensory filtering can arise from plastic changes in inhibitory circuits rather than static inhibition alone (Das et al., 2011; Kato et al., 2015; Lamiré et al., 2023). Panx1 is also implicated in the regulation of inhibitory synaptic transmission and excitation–inhibition balance in hippocampal circuits, where disruption of Panx1 signaling alters GABAergic transmission and network excitability (Ardiles et al., 2014; Flores-Muñoz et al., 2022; Illanes-González et al., 2025), providing a potential mechanism for the impaired stabilization of habituation observed in *panx1a^−/−^* larvae.

### Network-level coordination across oscillations and transient events

At the network level, experience-dependent plasticity is reflected in coordinated changes across oscillatory activity and transient population events. In wild-type larvae, habituation was associated with a reduction in gamma-band power in the ADL, a region functionally analogous to the mammalian hippocampus. Gamma oscillations arise from interactions between excitatory neurons and interneurons and are widely interpreted as a marker of local circuit synchronization (Colgin, 2016). The observed reduction in gamma activity is therefore consistent with a reorganization of local network dynamics following repeated stimulation. The absence of comparable modulation in *panx1a^−/−^* larvae indicate impaired adjustment of these coordinated activity patterns.

Changes were also evident at the level of cross-frequency interactions. In wild-type larvae, habituation reduced theta–gamma phase–amplitude coupling (PAC) in the ADL, consistent with a shift in network state. PAC reflects structured interactions between low-frequency phase and high-frequency amplitude and is thought to support the coordination of neural activity across temporal and spatial scales (Soulat et al., 2022). Recent work demonstrates that hippocampal theta–gamma PAC coordinates interactions between frontal and medial temporal lobe circuits and tracks working memory demand and behavioral performance (Daume et al., 2024). In this context, the reduction in PAC observed here is consistent with a transition from a highly coordinated, novelty-responsive state toward a more stable processing regime. This adaptation was absent in *panx1a^−/−^* larvae, suggesting reduced flexibility in experience-dependent network reconfiguration.

At the inter-regional level, coherence between the optic tectum and ADL increased in the theta band following habituation in wild-type larvae but was disrupted in mutants. This indicates that Panx1a contributes not only to local circuit dynamics but also to coordination across regions within the visual processing pathway. Together with the PAC findings, these results point to a role for Panx1a in supporting to organize activity across multiple temporal and spatial scales.

Experience-dependent effects were also evident in transient network events. SWR-like complexes were present in the developing pallium in vivo and exhibited selective modulation following habituation. Specifically, the duration of the SW component was reduced in wild-type larvae, whereas ripple-associated features, including duration and event frequency, remained unchanged. This dissociation indicates that experience-dependent plasticity preferentially affects the low-frequency component of these events at this developmental stage. In *panx1a^−/−^* larvae, this refinement of SW duration was attenuated, while overall event occurrence and ripple properties were preserved. These findings indicate that Panx1a is not required for the generation of SWR-like events but contributes to their temporal structuring.

Altogether, these observations suggest that experience-dependent plasticity in the developing brain operates through coordinated adjustments of both ongoing oscillatory activity and transient population events. Across measures, Panx1a loss is associated with reduced modulation of gamma activity, impaired PAC adaptation, disrupted inter-regional coherence, and diminished refinement of sharp-wave dynamics. Rather than reflecting a global disruption of activity, this pattern is consistent with a reduced capacity to reorganize network dynamics in response to experience.

### A unifying framework across scales

Taken together, our findings support a model in which Panx1a functions as a mediator of activity-dependent signaling across multiple levels of organization. Through ATP release and purinergic signaling, Panx1a may couple neural activity to intracellular pathways that regulate transcription, synaptic plasticity, and network coordination. Loss of Panx1a disrupts this coupling, leading to deficits that manifest at the behavioral, molecular, and electrophysiological levels. Rather than acting within a single pathway, Panx1a appears to shape how neural activity is translated into stable adaptations. In this framework, the impaired habituation observed in *panx1a^−/−^* larvae reflects a broader failure to stabilize experience-dependent changes over time.

### Energy and efficiency in habituation

Although we did not directly measure metabolic activity, the dual role of Panx1 in ATP release and purinergic signaling raises the possibility that it contributes to coupling neural activity with energetic regulation (G. S. O. Zoidl et al., 2025). Habituation has been proposed to reflect an optimization process in which neural responses are reduced as stimuli become predictable, balancing information processing with energetic cost. Our results are consistent with this framework, suggesting that Panx1a may participate in signaling pathways that constrain circuit activity during repeated stimulation, although direct tests of this hypothesis will be required.

### Limitations and future directions

General limitations should be considered. Transcriptional and co-activation measures remain correlational and do not establish causal relationships between brain regions or processes. Pharmacological manipulations may have indirect effects on circuit excitability, and developmental compensation in *panx1a^−/−^* larvae cannot be excluded. Additionally, the temporal dynamics of transcriptional responses may differ between genotypes, and our sampling window may not capture all relevant changes. Despite these limitations, the convergence of behavioral, molecular, pharmacological, and electrophysiological findings supports a model in which Panx1a is a key regulator of experience-dependent plasticity. By facilitating the coupling of neural activity to transcriptional and network-level responses, Panx1a contributes to the brain’s ability to filter repetitive stimuli and adapt to a changing sensory environment.

## Methods

### Zebrafish Husbandry

Tupfel long fin strains of zebrafish were used as wild-type (WT) (Tupfel-longfin - wildtype) larvae. The *panx1a^−/−^* mutant lines were generated from these Tupfel long-fin strains via TALEN technology (Safarian et al., 2020). The knockout lines were bred to at least the 3rd generation before testing on their larvae. The fishes were maintained at 28°C on a 14-hour light/10-hour dark cycle in a recirculation system (Aquaneering Inc., San Diego, CA).

### Behavioral testing of 6-dpf larvae

**Set up** - 6 days post fertilization (dpf) larvae were tested using a Zebrabox® behaviour recording system (ViewPoint Life Technology, Lyon, France; http://www.viewpoint.fr), and the Zebralab software (ViewPoint Life Technology, Lyon, France; http://www.viewpoint.fr). Activity quantization was used to record the response of larvae to the light/dark flashes during the training and testing blocks. For each block, larvae were subjected to 60 light flashes lasting 5 seconds, occurring at every 30 seconds. The larvae were trained for three training blocks separated by 30-minutes, and their response was tested after 2hrs and 24hrs to the same stimuli. Light controls were adjusted using a lightbox and were set to 30% visible light for the light-ON segment and 0% for the light-OFF segment. All experiments were performed at 28°C and were performed simultaneously at consistent times of the day to minimize the confounding factors that affect the larval behaviour, such as the circadian clock.

### Pharmacological assays

For pharmacological manipulations, 6-dpf larvae were incubated in E3 medium containing the indicated drug or vehicle control for 1 hr prior to behavioral training. The following compounds were used: AP5 (50 µM) (Wolman et al., 2011), MK801 (20 µM) (Zoodsma et al., 2020), Memantine (30 µM) (Best et al., 2008), DNQX (20 µM) (Bhandiwad et al., 2018), Diazepam (0.75 µM), Gabazine (10 µM) (Jadhav et al., 2025), Bicuculline (5 µM) (Lamiré et al., 2023), Glycine (1 mM), and Actinomycin D (10 µM). Drug concentrations were adapted from published literature for the duration of the visual habituation assay and were selected to avoid effects on kinematic performance or spontaneous initiation of light-evoked bends. Larvae were maintained in drug solution throughout training and testing, unless otherwise noted.

For behavioural assays, control and treated groups underwent the standard training protocol, followed by testing at 2 hr post-training. Habituation percent was calculated relative to baseline performance as described above. For each drug condition, both WT and *panx1a⁻/⁻* larvae were tested in parallel to enable direct genotype comparisons.

### Click chemistry in 6-pf larvae

Zebrafish larvae (6-dpf, WT) were incubated with 2.5mM 5-Ethydyl uridine (5-EU) in egg water and tested for habituation paradigm (3 training blocks, 2hr testing block). After testing 2hr response, the larvae were washed 5 times for 2-3 minutes with ice-cold egg water to anesthetize them and were fixed in cold PFA-fixative (4% PFA, 4% sucrose). The larvae were incubated overnight at 4°C. Next day, larvae were digested with 1 mg/mL Collagenase for 1 hr at room temperature and were then washed briefly in PBS-Tween, pH 7.4. They were immediately post-fixed for 20 minutes in PFA-sucrose at room temperature. The click-reaction was adjusted to 500uL in individual wells and contained 100mM CuSO4, 100mM BTTAA, 100mM sodium phosphate buffer and 10mM Alexa 594-azide. Larvae were stained for DAPI in PBS buffer overnight before mounting in 0.6% low-melt agarose.

### Hybridization chain reaction RNA-fluorescence in situ hybridization (HCR RNA-FISH)

Larvae were raised under standard conditions in egg water. When embryos reached 12 hours post-fertilization (hpf), egg water was replaced with egg water containing 0.003% of 1-phenyl 2-thiourea (PTU). Fresh egg water containing 0.003% PTU was replaced every 24 hours until the larvae reached 5 dpf. At 6-dpf, the larvae were treated for behavioural assay or left in dark; and fixed in 4% ice-cold paraformaldehyde (PFA). After 24 hours, the PFA was washed out with 1× PBS, and the samples were gradually dehydrated, permeabilized with methanol, and stored at −20°C for several days until HCR in-situ labelling was performed. Staining was performed according to the manufacturer’s protocol for whole-mount zebrafish larvae (Choi et al., 2018). Samples were separated into 5 larvae per well in a 24-well. plate. Rehydration steps were performed by washing for 5 minutes (mins) each in 75% methanol/PBST (1× PBS + 0.1% Tween-20), 50% methanol/PBST, 25% methanol/PBST, and 5 times with 100% PBST. The samples were permeabilized with 30 µg/ml proteinase K for 45 mins at room temperature (RT), followed by post-fixation with 4% PFA for 20 mins at RT, and 5 washes in PBST for 5 mins each. The samples were prehybridized in 500 µl of probe hybridization buffer (Molecular Instruments) for 30 mins at 37°C. Hybridization was performed by adding 2 pmol of each probe set to the hybridization buffer and incubating for 16 hours at37°C.

Probe sets for *panx1a* and *fosab* were purchased from and designed by Molecular Instruments using proprietary HCR methodology to detect and fluorescently label target RNA transcripts. To maximize targeting, the probe was designed against shared regions of known variants found on NCBI Gene and the Ensembl database. Each probe set consisted of 20 split-initiator probe pairs per target and utilized a B1 amplifier with a 546 nm fluorophore label. To remove excess probes, the samples were washed 4 times for 15 mins each with a wash buffer (Molecular Instruments) at 37°C, followed by 2 washes of 5 mins each with 5× SSCT (5× SSC + 0.1% Tween-20) at RT. Pre-amplification was performed by incubating the samples for 30 mins in an amplification buffer (Molecular Instruments) at RT. The fluorescently labelled hairpins (B2-488 for *fosab* and B1-594 for *panx1a*) were prepared by snap cooling: heating at 95°C for 90 seconds and then cooling to RT for 30 mins. The hairpin solution was prepared by adding 10 µl of the snap-cooled hairpins (3 µM stock concentration) to 500 µl of amplification buffer. The pre-amplification buffer was removed, and the samples were incubated in the hairpin solution for 16 hours at RT. Excess hairpins were washed three times with 5× SSCT for 20 mins each. Following HCR RNA-FISH, larvae were stained with DAPI (1:12,000) overnight at 4°C, followed by three 10-minute washes in 5× SSCT. The samples were then stored in 5× SSCT in the dark at 4°C until imaging. A negative control was included, where no probes were added but fluorescent hairpins were used. The larvae were raised under standard conditions, euthanized at 6 dpf, depigmented in peroxide and potassium hydroxide treatment, and processed for HCR in-situ labeling.

### Confocal image acquisition and analysis

A z-mold or an 8-teeth mold for zebrafish larvae was 3D-printed using 2% low melting agarose on glass-bottom culture dishes (MatTek, P35G-0-10C) (Geng and Peterson, 2021). The larvae were positioned in larva-shaped slots and embedded dorsal side down in 0.5% low melting agarose. Images were acquired on the Nikon A1R confocal system with 10x air or 20x water objectives using the 546 nm, 488 nm and 468 nm lasers. Image analysis was performed using FIJI (v2.9.0). The acquired z-stacks were mapped against the mapZebrain atlas (Kunst et al., 2019; Shainer et al., 2023) using the DAPI stain channel as a bridge (see reference brain mapping in materials and methods). Mean raw fluorescence values were then extracted from well-defined anatomical mapZebrain atlas regions.

### Reference brain mapping

We used the ANTs brain registration library (Avants et al., 2011) to map volumes against the mapZebrain atlas (Kunst et al., 2019; Shainer et al., 2023) following a previously established approach applied to another pannexin channel, Pannexin2 (Shanbhag et al., 2025). We first used Fiji to manually convert each acquired confocal volume into separate nrrd files, one for the DAPI and one for the *panx1a* channel/*fosab* channel. To then generate a warp transformation file across coordinate systems, we mapped DAPI volumes against the T_AVG_DAPI reference volume (downloaded from the mapZebrain atlas platform). We then applied the resulting transform to the *panx1a* probe/*fosab* probe channel. We used brain region masks (downloaded from the mapZebrain platform) to slice out the fluorescence from the mapped image stacks for further analysis of the *panx1a* or *fosab* expression levels within well-defined separate brain areas.

For all mappings, we used the following ANTs commands:

antsRegistration -v 1 -d 3 --float 1 --winsorize-image-intensities [0.005, 0.995] --use-histogram-
matching 0 -o <output.nrrd> --initial-moving-transform
[<DAPI_reference.nrrd>,<image_DAPI_channel.nrrd>,1] -t Rigid[0.1] -m
MI[<DAPI_reference.nrrd>,<image_DAPI_channel.nrrd>,1,32,Regular,0.5] -c
[1000×500×250×300,1e-8,10] -s 3×2×1×0 -f 8×4×2×1 -t Affine[0.1] -m
MI[<DAPI_reference.nrrd>,<image_DAPI_channel.nrrd>,1,32,Regular,0.5] -c
[200×200×200×100,1e-8,10] -s 3×2×1×0 -f 8×4×2×1 -t SyN[0.1,6,0] -m
CC[<DAPI_reference.nrrd>,<image_DAPI_channel.nrrd>,1,2] -c [200×100,1e-7,10] -s 4×3 -f
12×8 antsApplyTransforms -d 3 -v 0 -- float -n linear -i <image_panx2_channel.nrrd> -r
<DAPI_reference.nrrd> -o <image_panx2_channel_mapped.nrrd> -t <image_1Warp.nii.gz> -t <
image_0GenericAffine.mat>

### Fluorescence signal analysis post registration

Mean *fosab* signal intensities were quantified across brain regions in larval zebrafish under different training conditions and genotypes (WT and *panx1a−/−*). The dataset was filtered to include only those brain regions in WT fish that had signal data from both Naive and Trained groups. For these regions, a Wilcoxon rank-sum test (Mann-Whitney U test) was performed to identify significantly modulated regions (p < 0.05) between WT Naive and WT Trained groups. The heatmap displays the mean *fosab* signal intensity for each group (WT Naive, WT Trained, KO Naive, KO Trained) across these significant brain regions. The brain regions (columns) are ordered based on the effect size of training in KO larvae, calculated as the difference in mean signal between KO Trained and KO Naive groups. Each row in the heatmap corresponds to a group, and each column to a brain region. Signal intensity is represented using a custom diverging color palette ranging from dark green to bright red.

Mean fluorescence signal intensity from click-chemistry labeling was calculated across brain regions in WT and *panx1a^−/−^* larval zebrafish under Naive and Trained conditions. For each brain region, the mean signal in KO_Trained and KO_Naive groups was compared, and the top 50 brain regions were selected based on the largest training-related change (KO_Trained – KO_Naive). These regions were then plotted in a horizontal heatmap showing mean signals for all four experimental groups: WT_Naive, WT_Trained, KO_Naive, and KO_Trained. Rows in the heatmap represent groups, and columns represent individual brain regions. The brain regions are ordered from left to right by increasing training-related effect size in KO animals. A custom diverging color palette ranging from dark green to red was used to represent signal strength.

### In vivo electrophysiology

Published procedures were used to anesthetize 6-dpf zebrafish larvae for in vivo electrophysiology (Baraban, 2013; Safarian et al., 2020; Whyte-Fagundes et al., 2022). Zebrafish larvae at 6-dpf with or without 4hrs MPTP treatment were anesthetized with 0.3 mM Pancuronium bromide (Sigma‒Aldrich, Oakville, ON, Canada) for 2‒3 min until the touch response stopped. Anesthetized larvae were immobilized in freshly prepared 2% low-melting temperature agarose. A Leica S9E dissecting microscope (Leica Microsystems, Richmond Hill, ON, Canada) was used to orient the dorsal aspect of the larvae to the gel surface. The embedded larvae were placed on the upright stage of an Olympus BX51 fluorescence microscope (Olympus, Richmond Hill, ON, Canada). Larvae were submerged in 1 ml of egg mixture (E3; pH 7.2-7.4) applied topically to the agar. Under direct visual guidance, two glass microelectrodes (1.2 mM OD, approximately 1 µM tip diameter, 2–7 MΩ), backloaded with 2 M NaCl, were placed into the right optic tectum (OT) and the dorsolateral (ADL) region of the pallium. Both microelectrodes recorded local field potentials simultaneously via a Multiclamp 700B amplifier (Axon Instruments, San Jose, CA, USA). The voltage recordings were low-pass filtered at 1 kHz (−3 dB; eight-pole Bessel), high-pass filtered at 0.1 Hz, digitized at 10 kHz via a Digidata 1550A A/D interface, and stored on a PC running pClamp11 software (all from Axon Instruments). The basal activity was recorded for 10 minutes under Light-ON conditions (1000 lux), during which images of the electrode placement were taken for reference. For each fish, brain activity was normalized to its baseline activity to account for the biological variability of individual brains. In the treatment group, larvae were habituated and exposed to the short-term recall, followed by immediate electrophysiological recording. A temperature of 28°C was maintained during the experiments.

### Power spectral density and coherence detection analysis from continuous LFP recording

Power spectral density (PSD) and coherence estimation were performed using NeuroExplorer Version 4 (Nex Technologies, Colorado Springs, CO, 80906, USA). To calculate PSD the original data were split into 65536 data segments. Each interval in a rate histogram was detrended by linear regression, and subjected to Fourier transform (FFT) using Welch’s method of windowing (1, 6, 7). A raw power spectrum was generated from the FFTs. Normalized spectral densities for continuous variables were determined for the frequency bands theta (3–7 Hz) and gamma (35–45 Hz) and used for statistical analysis in GraphPad Prism VS10. All data within their respective frequency bands were averaged per animal for visualization and statistical analysis. To measure Coherence, as a function of frequency, between two time series with continuous variables digitized at the same frequency, the FFTs were calculated after detrending and applying Welch’s method of tapering. Confidence levels were calculated as described in Kattla and Lowery (Kattla & Lowery, 2010). Changes in both PSD and coherence between genotypes and training were analyzed for significance via Welch’s t-test in GraphPad Prism.

### Phase-Amplitude Coupling Analysis

The phase amplitude coupling (PAC) was calculated using the *pactools* code (Tour et al., 2017). The code provides calculations for several PAC methods. Here, a PAC was performed based on the Hilbert transformation and the generalized linear models (GLM) (Penny et al., 2008). The raw LFP recordings were downsampled to 1000Hz. A custom data loader was created to load in the “.abf” file format. The phase from one to 12 hertz (Hz) was calculated to determine how it modulates the amplitudes of beta and gamma oscillations (13 - 100Hz) for one-second timeframes. Following the PAC analysis using the GLM model, the results were averaged and visualized using comodulograms. The unitless beta-coefficients (PAC strength) were extracted from the comodulograms and compared between the different treatments and genotypes. Changes in PAC strength between baseline, treatment, and genotype were determined using a two-way ANOVA with Tukey’s multiple comparisons test.

### Sharp Wave Ripple Complex Analysis

LFP recordings were analyzed using a custom Python pipeline. Signals were imported, and processed channel-wise. Line noise at 60 Hz and its harmonics was removed with notch filters, after which the signal was bandpass-filtered from 1-1,000 Hz using a zero-phase Gaussian-windowed finite impulse response filter. Sharp-wave and ripple components were then isolated using zero-phase bandpass filters at 1-30 Hz and 120-220 Hz, respectively.

Event detection was performed on the root mean square (RMS) envelopes of the filtered signals, calculated in sliding windows of 30ms for sharp waves and 5ms for ripples. Baseline mean and standard deviation were estimated for each RMS trace using a two-component Gaussian mixture model fitted to the full recording; the lower-mean component was taken as baseline. Sharp waves were detected when SW RMS exceeded 6 standard deviations (SDs) above baseline, with start and end times defined at 4 (SDs). Ripple events were detected when ripple RMS exceeded 4 (SDs) above baseline, with start and end times defined at 3 (SDs). Events shorter than 25ms were discarded, and events separated by less than 150ms were merged. SWR complexes were defined as sharp-wave events overlapping in time with at least one ripple. Where multiple ripples overlapped a single sharp wave, the ripple with the largest RMS peak was assigned. SWR parameters were extracted and averaged per animal. Outliers were removed using a ROUT test (Q = 1%). Significance was calculated using a two-factor linear model testing the effects of genotype, treatment, and their interaction, with Tukey-adjusted post hoc comparisons of estimated marginal means. R packages ‘car’ and ‘emmeans’ were used for statistical analysis.

### Statistics and data reproducibility

Statistical analyses were performed using GraphPad Prism (v10.2.3) and R. Data are presented as mean ± SD. Brain morphometric measurements were compared using the Kolmogorov–Smirnov test. Behavioral datasets were analyzed using two-way ANOVA followed by Šídák’s multiple comparisons tests. Electrophysiological measures (PSD, coherence) were analyzed using Welch’s t-tests, while phase–amplitude coupling (PAC) was assessed using two-way ANOVA with Tukey’s post hoc comparisons. *fosab* signal differences between groups were evaluated using the Wilcoxon rank-sum test. For SWR analyses, significance was determined using a two-factor linear model followed by Tukey-adjusted post hoc comparisons. Normality and homogeneity of variance were assessed using Shapiro-Wilk and Levene’s tests where applicable. A P value < 0.05 was considered statistically significant. Full statistical details, including sample sizes and exact P values, are provided in the figure legends and Supplementary Table 1.

## Acknowledgements

We thank two members of York University Zebrafish Vivarium, Janet Fleites-Medina, and Veronica Scavo for outstanding zebrafish husbandry.

## Author contributions

Conceptualization, FN, GRZ; data analysis, FN, GSZ, AB; investigation, all authors; writing – original draft preparation, FN, GRZ; writing - review and editing, all authors; visualization, FN, GSZ; supervision, GRZ; project administration, GRZ; funding acquisition, GRZ, AB.

## Consent for publication

All authors have read and agreed to the published version of the manuscript.

## Ethics approval

All animal work was performed at York University’s zebrafish vivarium and in an S2 biosafety laboratory following the Canadian Council for Animal Care guidelines after approval of the study 797 protocol by the York University Animal Care Committee (GZ#2019-7-R2).

## Data availability

All data supporting the findings of this study are available from the corresponding author upon request.

## Materials & Correspondence

Correspondence and material requests should be addressed to <u>Fatema Nakhuda</u>, <u>Georg R. Zoidl</u>

---

**Table.**
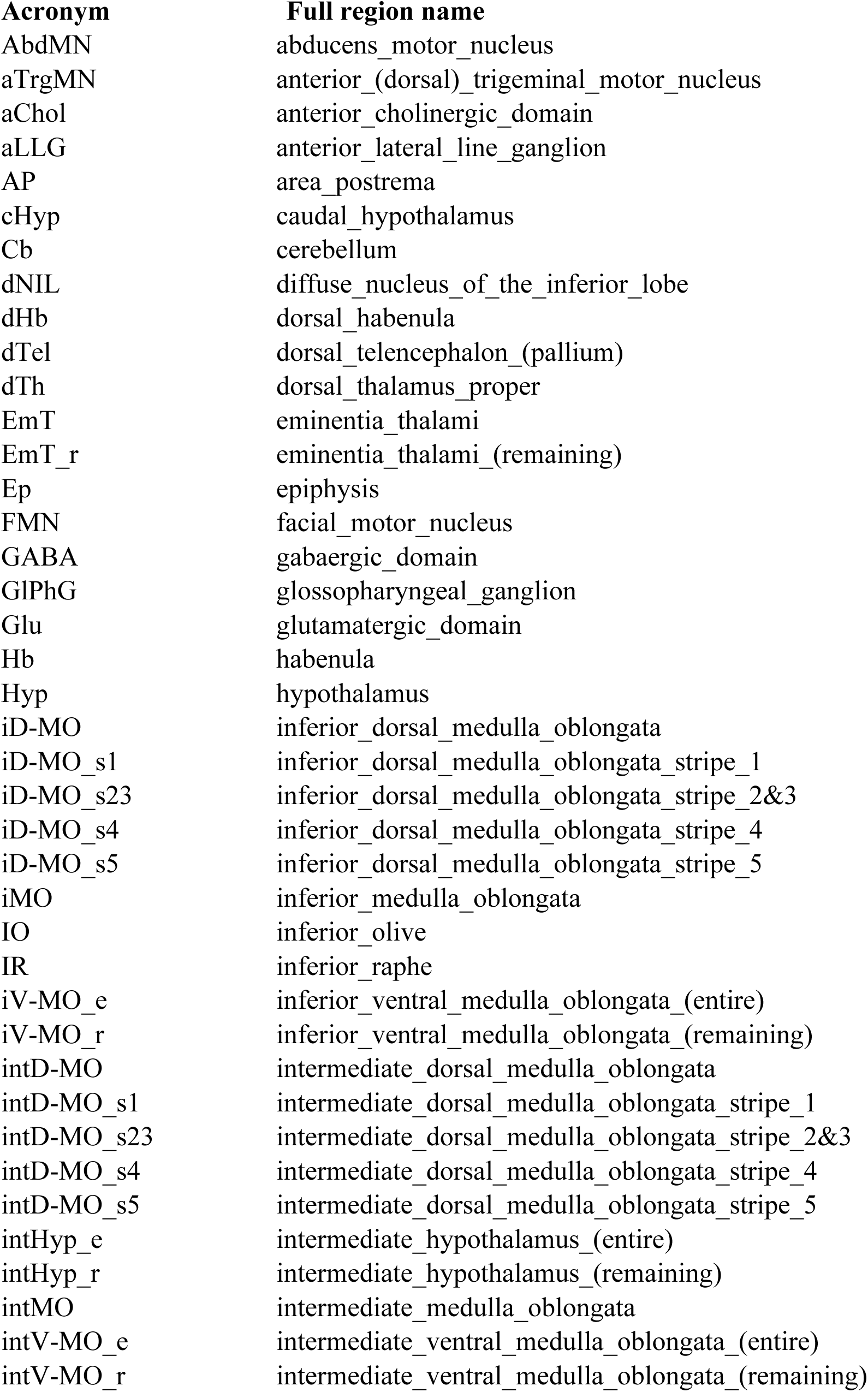
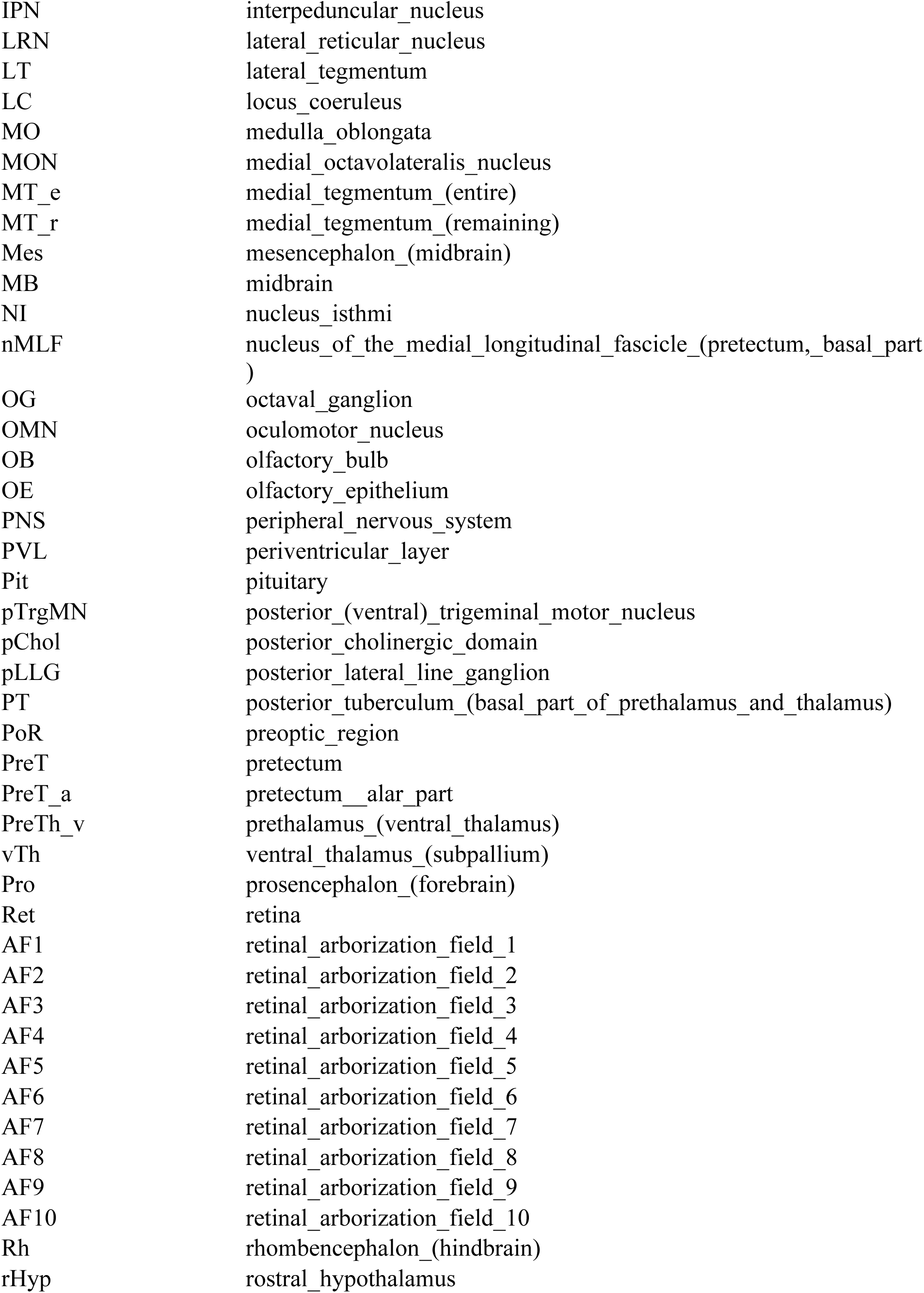
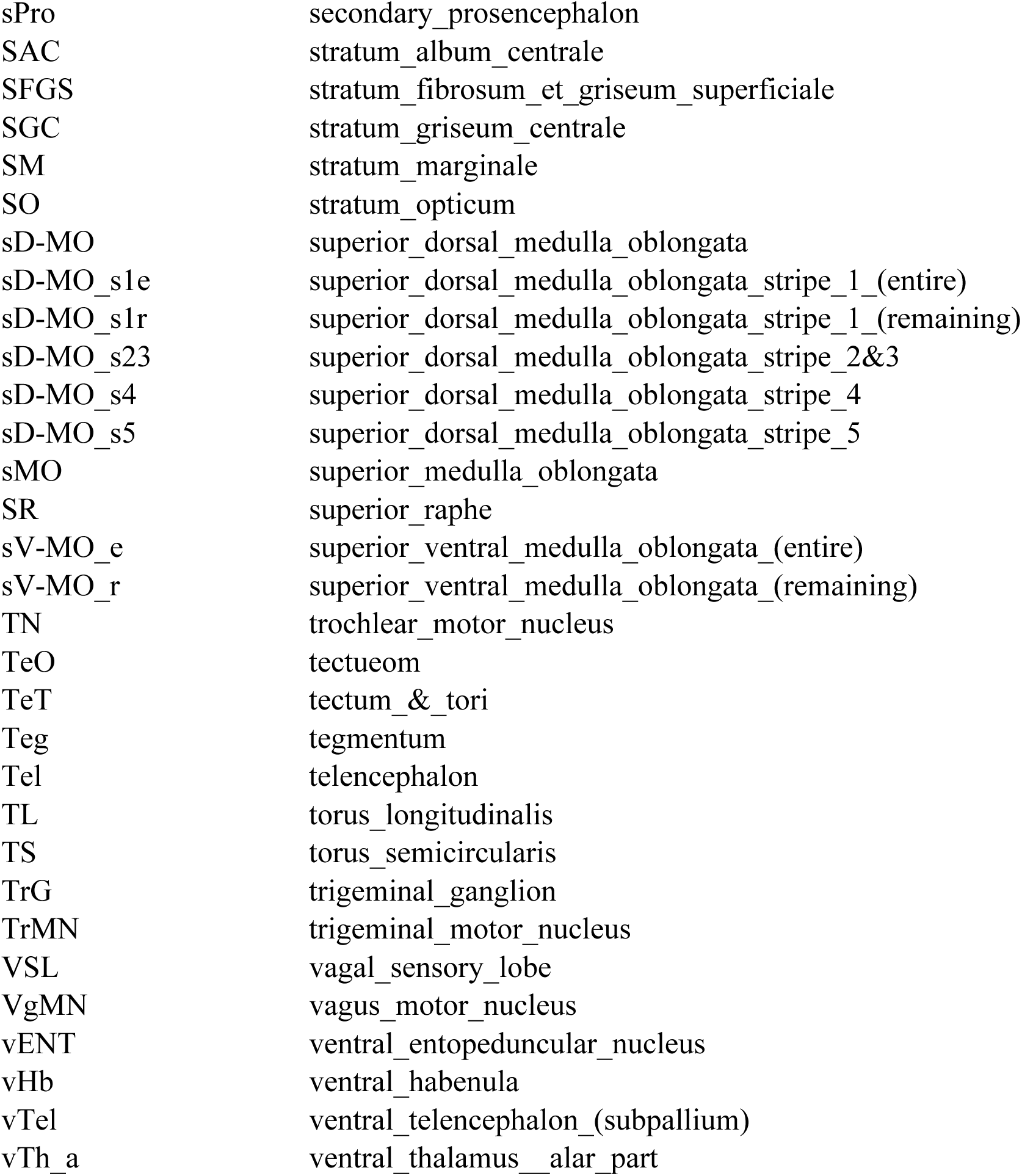
Region Acronyms in Figures 1 and 2.

## References

Adibi, M., Clifford, C. W. G., & Arabzadeh, E. (2013). Informational basis of sensory adaptation: Entropy and single-spike efficiency in rat barrel cortex. The Journal of Neuroscience: The Official Journal of the Society for Neuroscience, 33(37), 14921–14926. 10.1523/JNEUROSCI.1313-13.2013

Amilhon, B., Huh, C. Y. L., Manseau, F., Ducharme, G., Nichol, H., Adamantidis, A., & Williams, S. (2015). Parvalbumin Interneurons of Hippocampus Tune Population Activity at Theta Frequency. Neuron, 86(5), 1277–1289. 10.1016/j.neuron.2015.05.027

Ardiles, A. O., Flores-Muñoz, C., Toro-Ayala, G., Cárdenas, A. M., Palacios, A. G., Muñoz, P., Fuenzalida, M., Sáez, J. C., & Martínez, A. D. (2014). Pannexin 1 regulates bidirectional hippocampal synaptic plasticity in adult mice. Frontiers in Cellular Neuroscience, 8. 10.3389/fncel.2014.00326

Attwell, D., & Laughlin, S. B. (2001). An energy budget for signaling in the grey matter of the brain. Journal of Cerebral Blood Flow and Metabolism: Official Journal of the International Society of Cerebral Blood Flow and Metabolism, 21(10), 1133–1145. 10.1097/00004647-200110000-00001

Avants, B. B., Tustison, N. J., Song, G., Cook, P. A., Klein, A., & Gee, J. C. (2011). A reproducible evaluation of ANTs similarity metric performance in brain image registration. NeuroImage, 54(3), 2033–2044. 10.1016/j.neuroimage.2010.09.025

Bailey, C. H., Kandel, E. R., & Harris, K. M. (2015). Structural Components of Synaptic Plasticity and Memory Consolidation. Cold Spring Harbor Perspectives in Biology, 7(7), a021758. 10.1101/cshperspect.a021758

Bao, L., Locovei, S., & Dahl, G. (2004). Pannexin membrane channels are mechanosensitive conduits for ATP. FEBS Letters, 572(1–3), 65–68. 10.1016/j.febslet.2004.07.009

Baraban, S. C. (2013). Forebrain Electrophysiological Recording in Larval Zebrafish. Journal of Visualized Experiments : JoVE, (71), 50104. 10.3791/50104

Best, J. D., Berghmans, S., Hunt, J. J. F. G., Clarke, S. C., Fleming, A., Goldsmith, P., & Roach, A. G. (2008). Non-associative learning in larval zebrafish. Neuropsychopharmacology: Official Publication of the American College of Neuropsychopharmacology, 33(5), 1206–1215. 10.1038/sj.npp.1301489

Bhandiwad, A. A., Raible, D. W., Rubel, E. W., & Sisneros, J. A. (2018). Noise-Induced Hypersensitization of the Acoustic Startle Response in Larval Zebrafish. Journal of the Association for Research in Otolaryngology: JARO, 19(6), 741–752. 10.1007/s10162-018-00685-0

Blanco, I., Caccavano, A., Wu, J.-Y., Vicini, S., Glasgow, E., & Conant, K. (2024). Coupling of Sharp Wave Events between Zebrafish Hippocampal and Amygdala Homologs. The Journal of Neuroscience: The Official Journal of the Society for Neuroscience, 44(17), e1467232024. 10.1523/JNEUROSCI.1467-23.2024

Burgess, H. A., & Granato, M. (2007). Modulation of locomotor activity in larval zebrafish during light adaptation. Journal of Experimental Biology, 210(14), 2526–2539. 10.1242/jeb.003939

Burnstock, G., Fredholm, B. B., North, R. A., & Verkhratsky, A. (2010). The birth and postnatal development of purinergic signalling: Purinergic signalling: historic overview. Acta Physiologica, 199(2), 93–147. 10.1111/j.1748-1716.2010.02114.x

Buzsáki, G. (2015). Hippocampal sharp wave-ripple: A cognitive biomarker for episodic memory and planning. Hippocampus, 25(10), 1073–1188. 10.1002/hipo.22488

Buzsáki, G., & Wang, X.-J. (2012). Mechanisms of gamma oscillations. Annual Review of Neuroscience, 35, 203–225. 10.1146/annurev-neuro-062111-150444

Casillas Martinez, A., Wicki-Stordeur, L. E., Ariano, A. V., & Swayne, L. A. (n.d.). Dual role for pannexin 1 at synapses: Regulating functional and morphological plasticity. The Journal of Physiology, n/a(n/a). 10.1113/JP285228

Colgin, L. L. (2016). Rhythms of the hippocampal network. Nature Reviews. Neuroscience, 17(4), 239–249. 10.1038/nrn.2016.21

Cooke, S. F., Komorowski, R. W., Kaplan, E. S., Gavornik, J. P., & Bear, M. F. (2015). Visual recognition memory, manifested as long-term habituation, requires synaptic plasticity in V1. Nature Neuroscience, 18(2), 262–271. 10.1038/nn.3920

Dahl, G. (2015). ATP release through pannexon channels. Philosophical Transactions of the Royal Society B: Biological Sciences, 370(1672), 20140191. 10.1098/rstb.2014.0191

Das, S., Sadanandappa, M. K., Dervan, A., Larkin, A., Lee, J. A., Sudhakaran, I. P., Priya, R., Heidari, R., Holohan, E. E., Pimentel, A., Gandhi, A., Ito, K., Sanyal, S., Wang, J. W., Rodrigues, V., & Ramaswami, M. (2011). Plasticity of local GABAergic interneurons drives olfactory habituation. Proceedings of the National Academy of Sciences of the United States of America, 108(36), E646–654. 10.1073/pnas.1106411108

Daume, J., Kamiński, J., Schjetnan, A. G. P., Salimpour, Y., Khan, U., Kyzar, M., Reed, C. M., Anderson, W. S., Valiante, T. A., Mamelak, A. N., & Rutishauser, U. (2024). Control of working memory by phase–amplitude coupling of human hippocampal neurons. Nature, 629(8011), 393–401. 10.1038/s41586-024-07309-z

Esdin, J., Pearce, K., & Glanzman, D. L. (2010). Long-Term Habituation of the Gill-Withdrawal Reflex in Aplysia Requires Gene Transcription, Calcineurin and L-Type Voltage-Gated Calcium Channels. Frontiers in Behavioral Neuroscience, 4, 181. 10.3389/fnbeh.2010.00181

Ezzeddine, Y., & Glanzman, D. L. (2003). Prolonged Habituation of the Gill-Withdrawal Reflex in Aplysia Depends on Protein Synthesis, Protein Phosphatase Activity, and Postsynaptic Glutamate Receptors. The Journal of Neuroscience, 23(29), 9585–9594. 10.1523/JNEUROSCI.23-29-09585.2003

Flores-Muñoz, C., García-Rojas, F., Pérez, M. A., Santander, O., Mery, E., Ordenes, S., Illanes-González, J., López-Espíndola, D., González-Jamett, A. M., Fuenzalida, M., Martínez, A. D., & Ardiles, Á. O. (2022). The Long-Term Pannexin 1 Ablation Produces Structural and Functional Modifications in Hippocampal Neurons. Cells, 11(22), 3646. 10.3390/cells11223646

Fries, P. (2015). Rhythms for Cognition: Communication through Coherence. Neuron, 88(1), 220–235. 10.1016/j.neuron.2015.09.034

Gahtan, E., Tanger, P., & Baier, H. (2005). Visual Prey Capture in Larval Zebrafish Is Controlled by Identified Reticulospinal Neurons Downstream of the Tectum. Journal of Neuroscience, 25(40), 9294–9303. 10.1523/JNEUROSCI.2678-05.2005

Gajardo, I., Salazar, C. S., Lopez-Espíndola, D., Estay, C., Flores-Muñoz, C., Elgueta, C., Gonzalez-Jamett, A. M., Martínez, A. D., Muñoz, P., & Ardiles, Á. O. (2018). Lack of Pannexin 1 Alters Synaptic GluN2 Subunit Composition and Spatial Reversal Learning in Mice. Frontiers in Molecular Neuroscience, 11. https://www.frontiersin.org/article/10.3389/fnmol.2018.00114

García-Rojas, F., Flores-Muñoz, C., Santander, O., Solis, P., Martínez, A. D., Ardiles, Á. O., & Fuenzalida, M. (2023). Pannexin-1 Modulates Inhibitory Transmission and Hippocampal Synaptic Plasticity. Biomolecules, 13(6), 887. 10.3390/biom13060887

Greer, P. L., & Greenberg, M. E. (2008). From synapse to nucleus: Calcium-dependent gene transcription in the control of synapse development and function. Neuron, 59(6), 846–860. 10.1016/j.neuron.2008.09.002

Illanes-González, J., Flores-Muñoz, C., Vitureira, N., & Ardiles, Á. O. (2025). Pannexin 1 channels: A bridge between synaptic plasticity and learning and memory processes. Neuroscience and Biobehavioral Reviews, 174, 106173. 10.1016/j.neubiorev.2025.106173

Jadhav, M. P., Verma, S., & Thirumalai, V. (2025). Ionic conductances driving tonic firing in Purkinje neurons of larval zebrafish. The Journal of Physiology, 603(22), 7235–7253. 10.1113/JP286063

Kato, S., Kaplan, H. S., Schrödel, T., Skora, S., Lindsay, T. H., Yemini, E., Lockery, S., & Zimmer, M. (2015). Global brain dynamics embed the motor command sequence of Caenorhabditis elegans. Cell, 163(3), 656–669. 10.1016/j.cell.2015.09.034

Kattla, S., & Lowery, M. M. (2010). Fatigue related changes in electromyographic coherence between synergistic hand muscles. Experimental Brain Research, 202(1), 89–99. 10.1007/s00221-009-2110-0

Kendrick, K. M., Zhan, Y., Fischer, H., Nicol, A. U., Zhang, X., & Feng, J. (2011). Learning alters theta amplitude, theta-gamma coupling and neuronal synchronization in inferotemporal cortex. BMC Neuroscience, 12, 55. 10.1186/1471-2202-12-55

Khakh, B. S., & North, R. A. (2012). Neuromodulation by extracellular ATP and P2X receptors in the CNS. Neuron, 76(1), 51–69. 10.1016/j.neuron.2012.09.024

Koniaris, E., Drimala, P., Sotiriou, E., & Papatheodoropoulos, C. (2011). Different effects of zolpidem and diazepam on hippocampal sharp wave—Ripple activity *in vitro*. Neuroscience, 175, 224–234. 10.1016/j.neuroscience.2010.11.027

Kunst, M., Laurell, E., Mokayes, N., Kramer, A., Kubo, F., Fernandes, A. M., Förster, D., Dal Maschio, M., & Baier, H. (2019). A Cellular-Resolution Atlas of the Larval Zebrafish Brain. Neuron, 103(1), 21–38.e5. 10.1016/j.neuron.2019.04.034

Lamiré, L. A., Haesemeyer, M., Engert, F., Granato, M., & Randlett, O. (2023). Functional and pharmacological analyses of visual habituation learning in larval zebrafish. eLife, 12, RP84926. 10.7554/eLife.84926

Latini, S., & Pedata, F. (2001). Adenosine in the central nervous system: Release mechanisms and extracellular concentrations. Journal of Neurochemistry, 79(3), 463–484. 10.1046/j.1471-4159.2001.00607.x

Li, S., Bjelobaba, I., & Stojilkovic, S. S. (2018). Interactions of Pannexin1 Channels with Purinergic and NMDA Receptor Channels. Biochimica et Biophysica Acta, 1860(1), 166–173. 10.1016/j.bbamem.2017.03.025

Locovei, S., Bao, L., & Dahl, G. (2006). Pannexin 1 in erythrocytes: Function without a gap. Proceedings of the National Academy of Sciences, 103(20), 7655–7659. 10.1073/pnas.0601037103

Lyons, M. R., & West, A. E. (2011). Mechanisms of specificity in neuronal activity-regulated gene transcription. Progress in Neurobiology, 94(3), 259–295. 10.1016/j.pneurobio.2011.05.003

Ma, H., Khaled, H. G., Wang, X., Mandelberg, N. J., Cohen, S. M., He, X., & Tsien, R. W. (2023). Excitation–transcription coupling, neuronal gene expression and synaptic plasticity. Nature Reviews Neuroscience, 24(11), 672–692. 10.1038/s41583-023-00742-5

Malkani, S., & Rosen, J. B. (2000). Specific induction of early growth response gene 1 in the lateral nucleus of the amygdala following contextual fear conditioning in rats. Neuroscience, 97(4), 693–702. 10.1016/S0306-4522(00)00058-0

Mayford, M., Bach, M. E., Huang, Y.-Y., Wang, L., Hawkins, R. D., & Kandel, E. R. (1996). Control of Memory Formation Through Regulated Expression of a CaMKII Transgene. Science, 274(5293), 1678–1683. 10.1126/science.274.5293.1678

Mayhew, J., Graham, B. A., Biber, K., Nilsson, M., & Walker, F. R. (2018). Purinergic modulation of glutamate transmission: An expanding role in stress-linked neuropathology. Neuroscience & Biobehavioral Reviews, 93, 26–37. 10.1016/j.neubiorev.2018.06.023

Nicoletti, G., Bruzzone, M., Suweis, S., Maschio, M. D., & Busiello, D. M. (2025). Optimal information gain at the onset of habituation to repeated stimuli. eLife, 13. 10.7554/eLife.99767.2

North, R. A., & Verkhratsky, A. (2006). Purinergic transmission in the central nervous system. Pflugers Archiv: European Journal of Physiology, 452(5), 479–485. 10.1007/s00424-006-0060-y

Obot, P., Cibelli, A., Pan, J., Velíšek, L., Velíšková, J., & Scemes, E. (2024). Pannexin1 Mediates Early-Life Seizure-Induced Social Behavior Deficits. ASN Neuro, 16(1), 2371164. 10.1080/17590914.2024.2371164

Obot, P., Subah, G., Schonwald, A., Pan, J., Velíšek, L., Velíšková, J., Stanton, P. K., & Scemes, E. (2023). Astrocyte and Neuronal Panx1 Support Long-Term Reference Memory in Mice. ASN Neuro, 15, 17590914231184712. 10.1177/17590914231184712

Pankratov, Y., Lalo, U., Krishtal, O. A., & Verkhratsky, A. (2009). P2X receptors and synaptic plasticity. Neuroscience, 158(1), 137–148. 10.1016/j.neuroscience.2008.03.076

Patil, C. S., Li, H., Lavine, N. E., Shi, R., Bodalia, A., Siddiqui, T. J., & Jackson, M. F. (2022). ER-resident STIM1/2 couples Ca2+ entry by NMDA receptors to pannexin-1 activation. Proceedings of the National Academy of Sciences, 119(36), e2112870119. 10.1073/pnas.2112870119

Pellicano, M. P., Siciliano, F., & Sadile, A. G. (1993). NMDA receptors modulate long-term habituation to spatial novelty: Dose- and genotype-dependent differential effects of posttrial MK-801 and CPP in rats. Physiology & Behavior, 54(3), 563–568. 10.1016/0031-9384(93)90250-j

Penny, W. D., Duzel, E., Miller, K. J., & Ojemann, J. G. (2008). Testing for nested oscillation. Journal of Neuroscience Methods, 174(1), 50–61. 10.1016/j.jneumeth.2008.06.035

Pozo, K., & Goda, Y. (2010). Unraveling mechanisms of homeostatic synaptic plasticity. Neuron, 66(3), 337–351. 10.1016/j.neuron.2010.04.028

Prochnow, N., Abdulazim, A., Kurtenbach, S., Wildförster, V., Dvoriantchikova, G., Hanske, J., Petrasch-Parwez, E., Shestopalov, V. I., Dermietzel, R., Manahan-Vaughan, D., & Zoidl, G. (2012). Pannexin1 Stabilizes Synaptic Plasticity and Is Needed for Learning. PLoS ONE, 7(12), e51767. 10.1371/journal.pone.0051767

Randlett, O., Haesemeyer, M., Forkin, G., Shoenhard, H., Schier, A. F., Engert, F., & Granato, M. (2019). Distributed Plasticity Drives Visual Habituation Learning in Larval Zebrafish. Current Biology: CB, 29(8), 1337–1345.e4. 10.1016/j.cub.2019.02.039

Rangel-Sandoval, C., Soula, M., Li, W.-P., Castillo, P. E., & Hunt, D. L. (2024a). NMDAR-mediated activation of pannexin1 channels contributes to the detonator properties of hippocampal mossy fiber synapses. iScience, 27(5), 109681. 10.1016/j.isci.2024.109681

Rangel-Sandoval, C., Soula, M., Li, W.-P., Castillo, P. E., & Hunt, D. L. (2024b). NMDAR-mediated activation of pannexin1 channels contributes to the detonator properties of hippocampal mossy fiber synapses. iScience, 27(5), 109681. 10.1016/j.isci.2024.109681

Rankin, C. H., Abrams, T., Barry, R. J., Bhatnagar, S., Clayton, D. F., Colombo, J., Coppola, G., Geyer, M. A., Glanzman, D. L., Marsland, S., McSweeney, F. K., Wilson, D. A., Wu, C.-F., & Thompson, R. F. (2009). Habituation revisited: An updated and revised description of the behavioral characteristics of habituation. Neurobiology of Learning and Memory, 92(2), 135–138. 10.1016/j.nlm.2008.09.012

Ray, A., Zoidl, G., Weickert, S., Wahle, P., & Dermietzel, R. (2005). Site-specific and developmental expression of pannexin1 in the mouse nervous system. The European Journal of Neuroscience, 21(12), 3277–3290. 10.1111/j.1460-9568.2005.04139.x

Roberts, A. C., Pearce, K. C., Choe, R. C., Alzagatiti, J. B., Yeung, A. K., Bill, B. R., & Glanzman, D. L. (2016). Long-term habituation of the C-start escape response in zebrafish larvae. Neurobiology of Learning and Memory, 134 Pt B(Pt B), 360–368. 10.1016/j.nlm.2016.08.014

Roberts, A. C., Reichl, J., Song, M. Y., Dearinger, A. D., Moridzadeh, N., Lu, E. D., Pearce, K., Esdin, J., & Glanzman, D. L. (2011). Habituation of the C-start response in larval zebrafish exhibits several distinct phases and sensitivity to NMDA receptor blockade. PloS One, 6(12), e29132. 10.1371/journal.pone.0029132

Safarian, N., Whyte-Fagundes, P., Zoidl, C., Grigull, J., & Zoidl, G. (2020). Visuomotor deficiency in panx1a knockout zebrafish is linked to dopaminergic signaling. Scientific Reports, 10(1), 9538. 10.1038/s41598-020-66378-y

Sanchez-Arias, J. C., Candlish, R. C., van der Slagt, E., & Swayne, L. A. (2020). Pannexin 1 Regulates Dendritic Protrusion Dynamics in Immature Cortical Neurons. eNeuro, 7(4), ENEURO.0079-20.2020. 10.1523/ENEURO.0079-20.2020

Sanchez-Arias, J. C., Liu, M., Choi, C. S. W., Ebert, S. N., Brown, C. E., & Swayne, L. A. (2019). Pannexin 1 Regulates Network Ensembles and Dendritic Spine Development in Cortical Neurons. eNeuro, 6(3), ENEURO.0503-18.2019. 10.1523/ENEURO.0503-18.2019

Shainer, I., Kuehn, E., Laurell, E., Al Kassar, M., Mokayes, N., Sherman, S., Larsch, J., Kunst, M., & Baier, H. (2023). A single-cell resolution gene expression atlas of the larval zebrafish brain. Science Advances, 9(8), eade9909. 10.1126/sciadv.ade9909

Shanbhag, R., Zoidl, G. S. O., Nakhuda, F., Sabour, S., Naumann, H., Zoidl, C., Bahl, A., Tabatabaei, N., & Zoidl, G. R. (2025). Pannexin-2 deficiency disrupts visual pathways and leads to ocular defects in zebrafish. Biochimica et Biophysica Acta (BBA) - Molecular Basis of Disease, 1871(5), 167807. 10.1016/j.bbadis.2025.167807

Shigetomi, E., Sakai, K., & Koizumi, S. (2024). Extracellular ATP/adenosine dynamics in the brain and its role in health and disease. Frontiers in Cell and Developmental Biology, 11, 1343653. 10.3389/fcell.2023.1343653

Shimizu, E., Tang, Y.-P., Rampon, C., & Tsien, J. Z. (2000). NMDA Receptor-Dependent Synaptic Reinforcement as a Crucial Process for Memory Consolidation. Science, 290(5494), 1170–1174. 10.1126/science.290.5494.1170

Soulat, H., Stephen, E. P., Beck, A. M., & Purdon, P. L. (2022). State space methods for phase amplitude coupling analysis. Scientific Reports, 12(1), 15940. 10.1038/s41598-022-18475-3

Swayne, L. A., & Boyce, A. K. J. (2017). Regulation of Pannexin 1 Surface Expression by Extracellular ATP: Potential Implications for Nervous System Function in Health and Disease. Frontiers in Cellular Neuroscience, 11, 230. 10.3389/fncel.2017.00230

Thompson, R. F., & Spencer, W. A. (1966). Habituation: A model phenomenon for the study of neuronal substrates of behavior. Psychological Review, 73(1), 16–43. 10.1037/h0022681

Tort, A. B. L., Komorowski, R. W., Manns, J. R., Kopell, N. J., & Eichenbaum, H. (2009). Theta–gamma coupling increases during the learning of item–context associations. Proceedings of the National Academy of Sciences, 106(49), 20942–20947. 10.1073/pnas.0911331106

Tour, T. D. la, Tallot, L., Grabot, L., Doyère, V., Wassenhove, V. van, Grenier, Y., & Gramfort, A. (2017). Non-linear auto-regressive models for cross-frequency coupling in neural time series. PLOS Computational Biology, 13(12), e1005893. 10.1371/journal.pcbi.1005893

Turrigiano, G. G. (2008). The self-tuning neuron: Synaptic scaling of excitatory synapses. Cell, 135(3), 422–435. 10.1016/j.cell.2008.10.008

Tyssowski, K. M., DeStefino, N. R., Cho, J.-H., Dunn, C. J., Poston, R. G., Carty, C. E., Jones, R. D., Chang, S. M., Romeo, P., Wurzelmann, M. K., Ward, J. M., Andermann, M. L., Saha, R. N., Dudek, S. M., & Gray, J. M. (2018). Different Neuronal Activity Patterns Induce Different Gene Expression Programs. Neuron, 98(3), 530–546.e11. 10.1016/j.neuron.2018.04.001

Vogt, A., Hormuzdi, S. G., & Monyer, H. (2005). Pannexin1 and Pannexin2 expression in the developing and mature rat brain. Molecular Brain Research, 141(1), 113–120. 10.1016/j.molbrainres.2005.08.002

Weilinger, N. L., Lohman, A. W., Rakai, B. D., Ma, E. M. M., Bialecki, J., Maslieieva, V., Rilea, T., Bandet, M. V., Ikuta, N. T., Scott, L., Colicos, M. A., Teskey, G. C., Winship, I. R., & Thompson, R. J. (2016). Metabotropic NMDA receptor signaling couples Src family kinases to pannexin-1 during excitotoxicity. Nature Neuroscience, 19(3), 432–442. 10.1038/nn.4236

Whyte-Fagundes, P., Taskina, D., Safarian, N., Zoidl, C., Carlen, P. L., Donaldson, L. W., & Zoidl, G. R. (2022). Panx1 channels promote both anti- and pro-seizure-like activities in the zebrafish via p2rx7 receptors and ATP signaling. Communications Biology, 5(1), Article 1. 10.1038/s42003-022-03356-2

Wicki-Stordeur, L. E., & Swayne, L. A. (2013). Panx1 regulates neural stem and progenitor cell behaviours associated with cytoskeletal dynamics and interacts with multiple cytoskeletal elements. Cell Communication and Signaling, 11(1), 62. 10.1186/1478-811X-11-62

Wolman, M. A., Jain, R. A., Liss, L., & Granato, M. (2011). Chemical modulation of memory formation in larval zebrafish. Proceedings of the National Academy of Sciences, 108(37), 15468–15473. 10.1073/pnas.1107156108

Wulff, P., Ponomarenko, A. A., Bartos, M., Korotkova, T. M., Fuchs, E. C., Bähner, F., Both, M., Tort, A. B. L., Kopell, N. J., Wisden, W., & Monyer, H. (2009). Hippocampal theta rhythm and its coupling with gamma oscillations require fast inhibition onto parvalbumin-positive interneurons. Proceedings of the National Academy of Sciences of the United States of America, 106(9), 3561–3566. 10.1073/pnas.0813176106

Zoidl, G., Petrasch-Parwez, E., Ray, A., Meier, C., Bunse, S., Habbes, H.-W., Dahl, G., & Dermietzel, R. (2007). Localization of the pannexin1 protein at postsynaptic sites in the cerebral cortex and hippocampus. Neuroscience, 146(1), 9–16. 10.1016/j.neuroscience.2007.01.061

Zoidl, G. S. O., Safarian, N., Zoidl, C., Connor, S., & Zoidl, G. R. (2025). Panx1a modulates metabolic stress signaling and synaptic composition in the developing zebrafish brain. Cell and Tissue Research, 402(3), 217–242. 10.1007/s00441-025-04022-9

Zoodsma, J. D., Chan, K., Bhandiwad, A. A., Golann, D. R., Liu, G., Syed, S. A., Napoli, A. J., Burgess, H. A., Sirotkin, H. I., & Wollmuth, L. P. (2020). A Model to Study NMDA Receptors in Early Nervous System Development. The Journal of Neuroscience, 40(18), 3631–3645. 10.1523/JNEUROSCI.3025-19.2020

