## Supplemental Info. for "Multiscale regulation of experience-dependent plasticity by a Pannexin1 homolog in a developing vertebrate brain"

***Supplementary info.***

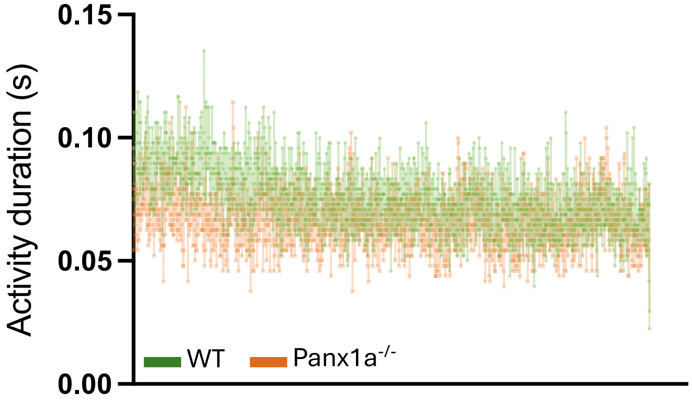

**Supplementary figure S1. Baseline locomotor activity** in the absence of stimuli, comparing WT and *panx1a*$^{-/-}$ larvae over a 30-minute period.

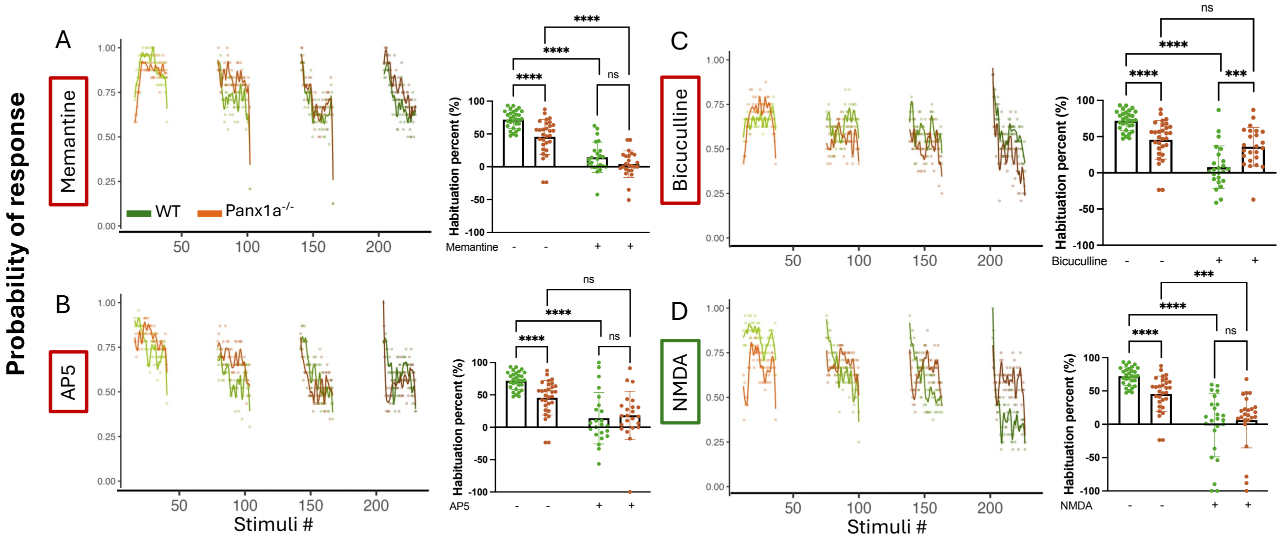

**Supplementary figure S2. Probability of response in habituation assay when different pharmacological agents are added. (A)** Non-competitive NMDA receptor antagonist that blocks glutamate binding at NMDA receptors, Memantine. **(B)** Selective competitive NMDA receptor antagonist AP5 **(C)** $GABA_A$ receptor antagonist that inhibits inhibitory signaling, Bicuculline **(D)** Enhancing NMDA activity using direct supplementation of NMDA.

**Table 1. Quantitative statistical summary of regional brain morphometrics and habituation responses across genotypes and pharmacological conditions.** Mean ± SD values, sample sizes (N), and corresponding statistical comparisons between WT and *panx1a-/-* larvae are reported. Brain region measurements (forebrain, tectum, corpus cerebelli, brainstem, and hindbrain) were analyzed using the Kolmogorov–Smirnov test. Habituation percentages under no drug and pharmacological treatments (Actinomycin D, DNQX, MK801, Glycine, Gabazine, and Diazepam) were analyzed using two-way ANOVA followed by Sidak's multiple comparisons test. Statistical significance is indicated as ns (not significant), $p < 0.05$ (*), $p < 0.01$ (**), $p < 0.001$ (***), and $p < 0.0001$ (****).

| Figure, Panel | Comparison | Genotype | N | Mean ± SD | p-value | P value summary |
|---|---|---|---|---|---|---|
| Figure 1, Panel E | Forebrain length | WT | 12 | 227.2 ± 19.15 | 0.2485 | ns |
| | | *panx1a*$^{-/-}$ | | 212.2 ± 20.38 | | |
| | Forebrain width | WT | | 183.2 ± 10.02 | <0.0001 | **** |
| | | *panx1a*$^{-/-}$ | | 161 ± 7.349 | | |
| | TeO length | WT | | 416.4 ± 33.14 | 0.0023 | ** |
| | | *panx1a*$^{-/-}$ | | 386.8 ± 15.11 | | |
| | TeO width | WT | | 198.5 ± 9.758 | 0.0097 | ** |
| | | *panx1a*$^{-/-}$ | | 178.5 ± 9.491 | | |
| | Corpus cerebelli Width | WT | | 351.1 ± 10.49 | 0.8475 | ns |
| | | *panx1a*$^{-/-}$ | | 346.1 ± 10.65 | | |
| | Brainstem Width | WT | | 235.6 ± 13.90 | 0.0996 | ns |
| | | *panx1a*$^{-/-}$ | | 223.1 ± 12.13 | | |
| | Hindbrain Length | WT | | 410.4 ± 19.64 | 0.2485 | ns |
| | | *panx1a*$^{-/-}$ | | 416.9 ± 7.641 | | |

| | | | | | | |
|---|---|---|---|---|---|---|
| Figure 1, Panel H | Block 1 | WT | 6 trials, 24 larvae per trial/ genotype | 100 ± 0.00 | | ns |
| | | *panx1a*$^{-/-}$ | | 100 ± 0.00 | | |
| | | WT | | 59.167 ± 1.169 | <0.0001 | **** |
| | | *panx1a*$^{-/-}$ | | 69.167 ± 1.169 | | |
| | Block 3 | WT | | 63.667 ± 1.633 | <0.0001 | **** |
| | | Panx1a-/- | | 69.333 ± 0.0816 | | |
| | Block 4 | *panx1a*$^{-/-}$ | | 81.833 ± 1.472 | <0.0001 | **** |
| | | Panx1a-/- | | 92.167 ± 0.753 | | |

| | | | | | | |
|---|---|---|---|---|---|---|
| Figure 1, Panel K | Habituation % - no drug | WT | 30 | 71.92 ± 13.609 | <0.0001 | **** |
| | | *panx1a*$^{-/-}$ | | 45.63 ± 26.643 | | |

| | | | | | | |
|---|---|---|---|---|---|---|
| | Habituation % - ActinomycinD | WT | | 46.96 ± 16.515 | <0.0001 | **** |
| | | *panx1a*$^{-/-}$ | | -5.217 ± 25.32 | | |
| | No Drug | WT | | 71.92 ± 13.609 | 0.0038 | ** |
| | ActinomycinD | | | 46.96 ± 16.515 | | |
| | No Drug | *panx1a*$^{-/-}$ | | 45.63 ± 26.643 | <0.0001 | **** |
| | ActinomycinD | | | -5.217 ± 25.32 | | |

| | | | | | | |
|---|---|---|---|---|---|---|
| Figure 2, Panel B | Habituation % - DNQX | WT | 30 | 45.421 ± 23.599 | 0.4491 | ns |
| | | *panx1a*$^{-/-}$ | | 50.430 ± 23.606 | | |
| | No Drug | WT | | 71.92 ± 13.609 | 0.0015 | ** |
| | DNQX | | | 45.421 ± 23.599 | | |
| | No Drug | *panx1a*$^{-/-}$ | | 45.63 ± 26.643 | 0.9999 | ns |
| | DNQX | | | 50.430 ± 23.606 | | |

| | | | | | | |
|---|---|---|---|---|---|---|
| Figure 2, Panel C | Habituation % - MK801 | WT | 30 | 18.472 ± 43.061 | <0.0001 | **** |
| | | *panx1a*$^{-/-}$ | | 45.729 ± 36.971 | | |
| | No Drug | WT | | 71.92 ± 13.609 | <0.0001 | **** |
| | MK801 | | | 18.472 ± 43.061 | | |
| | No Drug | *panx1a*$^{-/-}$ | | 45.63 ± 26.643 | 0.9999 | ns |
| | MK801 | | | 45.729 ± 36.971 | | |

| | | | | | | |
|---|---|---|---|---|---|---|
| Figure 2, Panel D | Habituation % - Glycine | WT | 30 | 24.595 ± 25.285 | 0.006 | ** |
| | | *panx1a*$^{-/-}$ | | 42.851 ± 24.337 | | |
| | No Drug | WT | | 71.92 ± 13.609 | <0.0001 | **** |
| | Glycine | | | 24.595 ± 25.285 | | |
| | No Drug | *panx1a*$^{-/-}$ | | 45.63 ± 26.643 | >0.9999 | ns |
| | Glycine | | | 42.851 ± 24.337 | | |

| | | | | | | |
|---|---|---|---|---|---|---|
| Figure 2, Panel E | Habituation % - Gabazine | WT | 30 | 28.169 ± 19.882 | 0.029 | ** |
| | | *panx1a*$^{-/-}$ | | 42.656 ± 16.048 | | |
| | No Drug | WT | | 71.92 ± 13.609 | <0.0001 | **** |
| | Gabazine | | | 28.169 ± 19.882 | | |
| | No Drug | *panx1a*$^{-/-}$ | | 45.63 ± 26.643 | >0.9999 | ns |
| | Gabazine | | | 42.656 ± 16.048 | | |

| | | | | | | |
|---|---|---|---|---|---|---|
| Figure 2, Panel F | Habituation % - Diazepam | WT | 30 | 46.596 ± 36.649 | <0.0001 | **** |
| | | *panx1a*$^{-/-}$ | | 19.547 ± 36.213 | | |

| | | | | | | |
|---|---|---|---|---|---|---|
| | No Drug | WT | | 71.92 ± 13.609 | 0.5265 | ns |
| | Diazepam | | | 46.596 ± 36.649 | | |
| | No Drug | *panx1a*$^{-/-}$ | | 45.63 ± 26.643 | 0.1225 | ns |
| | Diazepam | | | 19.547 ± 36.213 | | |
| Figure 4, Panel C | PSD - ADL Theta | WT - naïve | 10 | 0.4055 ± 0.2686 | 0.0025 | ** |
| | | *panx1a*$^{-/-}$ - naïve | 9 | 0.912 ± 0.3343 | | |
| | | WT - trained | 8 | 0.7866 ± 0.3688 | 0.4618 | ns |
| | | *panx1a*$^{-/-}$ - trained | 8 | 0.6488 ± 0.3597 | | |
| | Naïve | WT | 10 | 0.4055 ± 0.2686 | 0.03 | * |
| | Trained | | 8 | 0.7866 ± 0.3688 | | |
| | Naïve | *panx1a*$^{-/-}$ | 9 | 0.912 ± 0.3343 | 0.1412 | ns |
| | Trained | | 8 | 0.6488 ± 0.3597 | | |
| Figure 4, Panel D | PSD - TeO Theta | WT - naïve | 10 | 0.8658 ± 0.3726 | 0.7365 | ns |
| | | *panx1a*$^{-/-}$ - naïve | 9 | 0.9238 ± 0.3660 | | |
| | | WT - trained | 8 | 1.028 ± 0.3582 | 0.0749 | ns |
| | | *panx1a*$^{-/-}$ - trained | 8 | 0.7135 ± 0.2901 | | |
| | Naïve | WT | 10 | 0.8658 ± 0.3726 | 0.3626 | ns |
| | Trained | | 8 | 1.028 ± 0.3582 | | |
| | Naïve | *panx1a*$^{-/-}$ | 9 | 0.9238 ± 0.3660 | 0.2069 | ns |
| | Trained | | 8 | 0.7135 ± 0.2901 | | |
| Figure 4, Panel E | PSD - ADL Gamma | WT - naïve | 10 | 0.6069 ± 0.2751 | 0.0001 | *** |
| | | *panx1a*$^{-/-}$ - naïve | 9 | 0.04495 ± 0.0145 | | |
| | | WT - trained | 8 | 0.1531 ± 0.07546 | 0.0054 | ** |
| | | *panx1a*$^{-/-}$ - trained | 8 | 0.2903 ± 0.08988 | | |
| | Naïve | WT | 10 | 0.6069 ± 0.2751 | 0.0005 | *** |
| | Trained | | 8 | 0.1531 ± 0.07546 | | |
| | Naïve | *panx1a*$^{-/-}$ | 9 | 0.04495 ± 0.0145 | <0.0001 | **** |
| | Trained | | 8 | 0.2903 ± 0.08988 | | |
| | | WT - naïve | 10 | 0.1284 ± 0.06177 | 0.0012 | ** |

| | | | | | | |
|---|---|---|---|---|---|---|
| Figure 4, Panel F | PSD - TeO Gamma | *panx1a*$^{-/-}$ - naïve | 9 | 0.039 ± 0.02315 | | |
| | | WT - trained | 8 | 0.0552 ± 0.02164 | 0.0034 | ** |
| | | *panx1a*$^{-/-}$ - trained | 8 | 0.1935 ± 0.09176 | | |
| | Naïve | WT | 10 | 0.1284 ± 0.06177 | 0.0047 | ** |
| | Trained | | 8 | 0.0552 ± 0.02164 | | |
| | Naïve | *panx1a*$^{-/-}$ | 9 | 0.039 ± 0.02315 | 0.0018 | ** |
| | Trained | | 8 | 0.1935 ± 0.09176 | | |
| Figure 4, Panel H | PAC - ADL Theta Gamma PAC | WT - naïve | 10 | 0.04564 ± 0.02216 | <0.0001 | **** |
| | | *panx1a*$^{-/-}$ - naïve | 9 | 0.008512 ± 0.003888 | | |
| | | WT - trained | 8 | 0.01491 ± 0.005964 | <0.0001 | **** |
| | | *panx1a*$^{-/-}$ - trained | 8 | 0.008841 ± 0.003937 | | |
| | Naïve | WT | 10 | 0.04564 ± 0.02216 | <0.0001 | **** |
| | Trained | | 8 | 0.01491 ± 0.005964 | | |
| | Naïve | *panx1a*$^{-/-}$ | 9 | 0.008512 ± 0.003888 | 0.9412 | ns |
| | Trained | | 8 | 0.008841 ± 0.003937 | | |
| Figure 4, Panel J | PAC - TeO Theta Gamma PAC | WT - naïve | 10 | 0.02848 ± 0.007242 | <0.0001 | **** |
| | | *panx1a*$^{-/-}$ - naïve | 9 | 0.01714 ± 0.001180 | | |
| | | WT - trained | 8 | 0.05068 ± 0.007082 | <0.0002 | **** |
| | | *panx1a*$^{-/-}$ - trained | 8 | 0.02710 ± 0.005921 | | |
| | Naïve | WT | 10 | 0.02848 ± 0.007242 | <0.0003 | **** |
| | Trained | | 8 | 0.05068 ± 0.007082 | | |
| | Naïve | *panx1a*$^{-/-}$ | 9 | 0.01714 ± 0.001180 | <0.0004 | **** |
| | Trained | | 8 | 0.02710 ± 0.005921 | | |
| Figure 4, Panel L | Coherence - Theta | WT - naïve | 10 | 0.7157 ± 0.1579 | 0.5396 | ns |
| | | *panx1a*$^{-/-}$ - naïve | 9 | 0.7693 ± 0.1938 | | |
| | | WT - trained | 8 | 0.8638 ± 0.05304 | <0.0001 | **** |
| | | *panx1a*$^{-/-}$ - trained | 8 | 0.6323 ± 0.09000 | | |
| | Naïve | WT | 10 | 0.7157 ± 0.1579 | 0.0348 | * |

| | | | | | | |
|---|---|---|---|---|---|---|
| | Trained | | 8 | 0.8638 ± 0.05304 | | |
| | Naïve | | 9 | 0.7693 ± 0.1938 | 0.0822 | ns |
| | Trained | *panx1a*$^{-/-}$ | 8 | 0.6323 ± 0.09000 | | |
| Figure 4, Panel M | Coherance - Gamma | WT - naïve | 10 | 0.5229 ± 0.2332 | 0.2231 | ns |
| | | *panx1a*$^{-/-}$ - naïve | 9 | 0.3995 ± 0.1469 | | |
| | | WT - trained | 8 | 0.5290 ± 0.1373 | 0.7873 | ns |
| | | *panx1a*$^{-/-}$ - trained | 8 | 0.5114 ± 0.1177 | | |
| | Naïve | | 10 | 0.5229 ± 0.2332 | 0.9507 | ns |
| | Trained | WT | 8 | 0.5290 ± 0.1373 | | |
| | Naïve | | 9 | 0.3995 ± 0.1469 | 0.1022 | ns |
| | Trained | *panx1a*$^{-/-}$ | 8 | 0.5114 ± 0.1177 | | |
| Figure 5, Panel | SW Duration | WT - naïve | 18 | 241.46 ± 92.22 | 0.98 | ns |
| | | *panx1a*$^{-/-}$ - naïve | 16 | 231.78 ± 70.18 | | |
| | | WT - trained | 13 | 143.06 ± 55.05 | 0.21 | ns |
| | | *panx1a*$^{-/-}$ - trained | 13 | 203.2 ± 84.77 | | |
| | Naïve | | 18 | 241.46 ± 92.22 | 0.0055 | ** |
| | Trained | WT | 13 | 143.06 ± 55.05 | | |
| | Naïve | | 16 | 231.78 ± 70.18 | 0.76 | ns |
| | Trained | *panx1a*$^{-/-}$ | 13 | 203.2 ± 84.77 | | |
| Figure 5, Panel | Total SW number | WT - naïve | 18 | 499.44 ± 143.01 | 0.72 | ns |
| | | *panx1a*$^{-/-}$ - naïve | 16 | 556 ± 97.11 | | |
| | | WT - trained | 14 | 483.29 ± 156.95 | 0.9 | ns |
| | | *panx1a*$^{-/-}$ - trained | 13 | 446.38 ± 154.14 | | |
| | Naïve | | 18 | 499.44 ± 143.01 | 0.99 | ns |
| | Trained | WT | 13 | 483.29 ± 156.95 | | |
| | Naïve | | 16 | 556 ± 97.11 | 0.2 | ns |
| | Trained | *panx1a*$^{-/-}$ | 13 | 446.38 ± 154.14 | | |
| Figure 5, Panel | Ripple Duration | WT - naïve | 15 | 36.13 ± 5.66 | 0.7002 | ns |
| | | *panx1a*$^{-/-}$ - naïve | 16 | 38.23 ± 8.81 | | |

| | | | | | | |
|---|---|---|---|---|---|---|
| | | WT - trained | 14 | 38.34 ± 8.70 | 0.527 | ns |
| | | *panx1a*$^{-/-}$ - trained | 12 | 34.75 ± 5.32 | | |
| | Naïve | WT | 15 | 36.13 ± 5.66 | 0.8886 | ns |
| | Trained | | 14 | 38.34 ± 8.70 | | |
| | Naïve | *panx1a*$^{-/-}$ | 16 | 38.23 ± 8.81 | 0.3376 | ns |
| | Trained | | 12 | 34.75 ± 5.32 | | |
| Figure 5, Panel | Ripple Duration | WT - naïve | 17 | 87.47 ± 11.90 | 0.999 | ns |
| | | *panx1a*$^{-/-}$ - naïve | 16 | 87.94 ± 10.60 | | |
| | | WT - trained | 14 | 81.36 ± 21.90 | 0.678 | ns |
| | | *panx1a*$^{-/-}$ - trained | 13 | 88.77 ± 17.26 | | |
| | Naïve | WT | 17 | 87.47 ± 11.90 | 0.7557 | ns |
| | Trained | | 14 | 81.36 ± 21.90 | | |
| | Naïve | *panx1a*$^{-/-}$ | 16 | 87.94 ± 10.60 | 0.9998 | ns |
| | Trained | | 13 | 88.77 ± 17.26 | | |

**Supplementary Table 2.** ***Panx1a*** **RNA FISH expression mapping acronyms, as shown in Figure 1B.**

| **Acronym** | **Full region name** |
|---|---|
| AbdMN | abducens_motor_nucleus |
| aTrgMN | anterior_(dorsal)_trigeminal_motor_nucleus |
| aChol | anterior_cholinergic_domain |
| aLLG | anterior_lateral_line_ganglion |
| AP | area_postrema |
| cHyp | caudal_hypothalamus |
| Cb | cerebellum |
| dNIL | diffuse_nucleus_of_the_inferior_lobe |
| dHb | dorsal_habenula |
| dTel | dorsal_telencephalon_(pallium) |
| dTh | dorsal_thalamus_proper |
| EmT | eminentia_thalami |
| EmT_r | eminentia_thalami_(remaining) |
| Ep | epiphysis |
| FMN | facial_motor_nucleus |
| GABA | gabaergic_domain |
| GlPhG | glossopharyngeal_ganglion |
| Glu | glutamatergic_domain |
| Hb | habenula |
| Hyp | hypothalamus |
| iD-MO | inferior_dorsal_medulla_oblongata |
| iD-MO_s1 | inferior_dorsal_medulla_oblongata_stripe_1 |
| iD-MO_s23 | inferior_dorsal_medulla_oblongata_stripe_2&3 |
| iD-MO_s4 | inferior_dorsal_medulla_oblongata_stripe_4 |
| iD-MO_s5 | inferior_dorsal_medulla_oblongata_stripe_5 |
| iMO | inferior_medulla_oblongata |
| IO | inferior_olive |
| IR | inferior_raphe |
| iV-MO_e | inferior_ventral_medulla_oblongata_(entire) |
| iV-MO_r | inferior_ventral_medulla_oblongata_(remaining) |
| intD-MO | intermediate_dorsal_medulla_oblongata |
| intD-MO_s1 | intermediate_dorsal_medulla_oblongata_stripe_1 |
| intD-MO_s23 | intermediate_dorsal_medulla_oblongata_stripe_2&3 |
| intD-MO_s4 | intermediate_dorsal_medulla_oblongata_stripe_4 |
| intD-MO_s5 | intermediate_dorsal_medulla_oblongata_stripe_5 |
| intHyp_e | intermediate_hypothalamus_(entire) |
| intHyp_r | intermediate_hypothalamus_(remaining) |
| intMO | intermediate_medulla_oblongata |
| intV-MO_e | intermediate_ventral_medulla_oblongata_(entire) |

| | |
|---|---|
| intV-MO_r | intermediate_ventral_medulla_oblongata_(remaining) |
| IPN | interpeduncular_nucleus |
| LRN | lateral_reticular_nucleus |
| LT | lateral_tegmentum |
| LC | locus_coeruleus |
| MO | medulla_oblongata |
| MON | medial_octavolateralis_nucleus |
| MT_e | medial_tegmentum_(entire) |
| MT_r | medial_tegmentum_(remaining) |
| Mes | mesencephalon_(midbrain) |
| MB | midbrain |
| NI | nucleus_isthmi |
| nMLF | nucleus_of_the_medial_longitudinal_fascicle_(pretectum,_basal_part) |
| OG | octaval_ganglion |
| OMN | oculomotor_nucleus |
| OB | olfactory_bulb |
| OE | olfactory_epithelium |
| PNS | peripheral_nervous_system |
| PVL | periventricular_layer |
| Pit | pituitary |
| pTrgMN | posterior_(ventral)_trigeminal_motor_nucleus |
| pChol | posterior_cholinergic_domain |
| pLLG | posterior_lateral_line_ganglion |
| PT | posterior_tuberculum_(basal_part_of_prethalamus_and_thalamus) |
| PoR | preoptic_region |
| PreT | pretectum |
| PreT_a | pretectum__alar_part |
| PreTh_v | prethalamus_(ventral_thalamus) |
| vTh | ventral_thalamus_(subpallium) |
| Pro | prosencephalon_(forebrain) |
| Ret | retina |
| AF1 | retinal_arborization_field_1 |
| AF2 | retinal_arborization_field_2 |
| AF3 | retinal_arborization_field_3 |
| AF4 | retinal_arborization_field_4 |
| AF5 | retinal_arborization_field_5 |
| AF6 | retinal_arborization_field_6 |
| AF7 | retinal_arborization_field_7 |
| AF8 | retinal_arborization_field_8 |
| AF9 | retinal_arborization_field_9 |
| AF10 | retinal_arborization_field_10 |
| Rh | rhombencephalon_(hindbrain) |

| | |
|---|---|
| rHyp | rostral_hypothalamus |
| sPro | secondary_prosencephalon |
| SAC | stratum_album_centrale |
| SFGS | stratum_fibrosum_et_griseum_superficiale |
| SGC | stratum_griseum_centrale |
| SM | stratum_marginale |
| SO | stratum_opticum |
| sD-MO | superior_dorsal_medulla_oblongata |
| sD-MO_s1e | superior_dorsal_medulla_oblongata_stripe_1_(entire) |
| sD-MO_s1r | superior_dorsal_medulla_oblongata_stripe_1_(remaining) |
| sD-MO_s23 | superior_dorsal_medulla_oblongata_stripe_2&3 |
| sD-MO_s4 | superior_dorsal_medulla_oblongata_stripe_4 |
| sD-MO_s5 | superior_dorsal_medulla_oblongata_stripe_5 |
| sMO | superior_medulla_oblongata |
| SR | superior_raphe |
| sV-MO_e | superior_ventral_medulla_oblongata_(entire) |
| sV-MO_r | superior_ventral_medulla_oblongata_(remaining) |
| TN | trochlear_motor_nucleus |
| TeO | tectueom |
| TeT | tectum_&_tori |
| Teg | tegmentum |
| Tel | telencephalon |
| TL | torus_longitudinalis |
| TS | torus_semicircularis |
| TrG | trigeminal_ganglion |
| TrMN | trigeminal_motor_nucleus |
| VSL | vagal_sensory_lobe |
| VgMN | vagus_motor_nucleus |
| vENT | ventral_entopeduncular_nucleus |
| vHb | ventral_habenula |
| vTel | ventral_telencephalon_(subpallium) |
| vTh_a | ventral_thalamus__alar_part |

**Supplementary Table 3. 5-EU labelling expression values corresponding to panel D in Figure 2. Top 50 filtered based on fluorescent intensity values.**

| Brain Region | WT_Naive | WT_Trained | KO_Naive | KO_Trained |
|---|---|---|---|---|
| SO | 1641.417 | 1625.083 | 2899.25 | 1535.417 |
| SM | 1701.917 | 1671.083 | 2956.25 | 1606.167 |
| iD-MO | 1339.167 | 1336.25 | 2469.95 | 1123.667 |
| iD-MO_s4 | 1354.25 | 1313.1 | 2573.6 | 1233.417 |
| iD-MO_s23 | 1383.05 | 1308.625 | 2368.2 | 1071.333 |
| Cb | 1690.75 | 1647.25 | 2689 | 1435.75 |
| SFGS | 1482.917 | 1462.25 | 2591.083 | 1379 |
| iD-MO_s5 | 1452 | 1332.167 | 2399.625 | 1199.917 |
| MON | 1569.583 | 1513.667 | 2503.75 | 1326.583 |
| intD-MO_s4 | 1296.833 | 1268.25 | 2209.583 | 1043.583 |
| TeO | 1548.417 | 1531.417 | 2571.583 | 1421.083 |
| Ep | 2132.75 | 2127.583 | 3165.25 | 2014.833 |
| PVL | 1617 | 1600.333 | 2614.75 | 1467.75 |
| tectal_neuropil | 1445.167 | 1422.75 | 2495.25 | 1348.5 |
| AF10 | 1447.75 | 1425.083 | 2496.25 | 1351.75 |
| TeT | 1506.75 | 1491.917 | 2500.417 | 1376.667 |
| SFGS/SGC | 1342.25 | 1326.667 | 2348.083 | 1241.333 |
| MB | 1452.5 | 1438.25 | 2415.75 | 1315.75 |
| Mes | 1452.5 | 1438.25 | 2415.75 | 1315.75 |
| AF7 | 1175.583 | 1180.333 | 2080 | 984.4167 |
| iD-MO_s1 | 1145.375 | 1125.875 | 1954.25 | 865.75 |
| Hb | 1796.583 | 1791.583 | 2718.5 | 1630.25 |
| vHb | 1743.167 | 1743 | 2640.083 | 1552.333 |
| dHb | 1856.833 | 1846.417 | 2808.917 | 1726.417 |
| intD-MO | 1268.083 | 1216.083 | 2064 | 985.75 |
| sD-MO_s4 | 1433.333 | 1369.667 | 2201.333 | 1123.75 |
| intD-MO_s5 | 1347.958 | 1264 | 2074.458 | 1005.25 |
| SGC | 1263.25 | 1246.25 | 2223.792 | 1156.75 |
| intD-MO_s1 | 1225.833 | 1172.833 | 2023.083 | 959.875 |
| sD-MO_s5 | 1452.917 | 1374.75 | 2161.583 | 1109.708 |
| TL | 1884.833 | 1888.667 | 2849.667 | 1799.333 |
| intD-MO_s23 | 1207.167 | 1161.583 | 1999 | 954.1667 |
| OB | 1364.667 | 1370.667 | 2280.917 | 1241 |
| sac/spv | 1400.583 | 1378.75 | 2339.167 | 1303.083 |
| SAC | 1261.417 | 1239.25 | 2191.25 | 1157.083 |
| sD-MO | 1331.667 | 1304.917 | 2152.75 | 1120.583 |
| dTel | 1400.083 | 1394.75 | 2299.667 | 1268.833 |
| PreT_a | 1230.417 | 1230.417 | 2089.583 | 1061.167 |
| AF8 | 1145.333 | 1156.417 | 1947.375 | 934.5833 |

| | | | | |
|---|---|---|---|---|
| AF9 | 1065.083 | 1071.5 | 1895.833 | 886.5833 |
| PreT | 1198.417 | 1198.25 | 2037.667 | 1029.25 |
| sD-MO_s23 | 1297 | 1247.5 | 2028.667 | 1045.5 |
| FMN | 1067.708 | 1026.083 | 1745.583 | 766.625 |
| aTrgMN | 1162.75 | 1125.583 | 1807.708 | 829 |
| sD-MO_s1r | 1208.75 | 1166.167 | 1960.583 | 990 |
| sD-MO_s1e | 1192.667 | 1151.917 | 1945.083 | 977.25 |
| EmT_r | 1269.167 | 1284.792 | 2051.583 | 1096 |
| Tel | 1230.417 | 1236.417 | 2066.25 | 1113.417 |
| MT_r | 1126.667 | 1126.083 | 1928.75 | 976.3333 |
| Rh | 1060.167 | 1058.5 | 1738.417 | 812.75 |

**Supplementary Table 4. Fosab labelling expression values corresponding to panel G in Figure 2.**

| **Brain_Region** | **WT_Naive** | **WT_Trained** | **KO_Naive** | **KO_Trained** |
|---|---|---|---|---|
| AP | 966.8125 | 1436.5 | 1624 | 1000.375 |
| SR | 631 | 781.0714 | 906.0714 | 806.0833333 |
| LC | 677.25 | 828.6429 | 899.5357 | 811.0833333 |
| AF8 | 984.2917 | 1345.5 | 1138.036 | 1089.583333 |
| LT | 823.9167 | 989.3571 | 953.4286 | 915.3333333 |
| aChol | 761.1667 | 950.9286 | 940.3571 | 917.75 |
| GlPhG | 195.8 | 325.6 | 303.4167 | 293.625 |
| FMN | 914.8333 | 1123.429 | 969.9286 | 969.5 |
| Teg | 821.5833 | 1057.5 | 955.0714 | 965.5833333 |
| intD-MO_s23 | 858.6667 | 1097.5 | 891.1429 | 905.9166667 |
| AF3 | 772.3333 | 1049.679 | 1011.583 | 1036.625 |
| TN | 777.625 | 1023.5 | 907.1071 | 940.0833333 |
| aTrgMN | 677 | 902.25 | 897.1429 | 939.4166667 |
| intD-MO_s1 | 827.8333 | 1084.857 | 929.2143 | 971.5 |
| PT | 693.8333 | 1018.429 | 863.1429 | 907.1666667 |
| MT_e | 822.8333 | 1112.357 | 953.1429 | 1001.916667 |
| sD-MO_s1e | 772.0833 | 997.2143 | 895.3571 | 947.6666667 |
| MB | 904.5 | 1138.357 | 938.0714 | 991.3333333 |
| Mes | 904.5 | 1138.357 | 938.0714 | 991.3333333 |
| intD-MO | 894.5833 | 1117.929 | 892.9286 | 946.3333333 |
| sD-MO_s1r | 771.75 | 996.7857 | 894.5 | 948.4166667 |
| sD-MO_s23 | 805.8333 | 1025 | 895.5714 | 950.25 |
| MT_r | 831.25 | 1136.286 | 952.0714 | 1014.75 |
| TeO | 936.1667 | 1188.643 | 942.6429 | 1007 |
| TeT | 916.5833 | 1152 | 930.7143 | 996.3333333 |
| AF9 | 1246.5 | 1565.5 | 1275.357 | 1349.583333 |
| intD-MO_s4 | 919 | 1159.357 | 893.5 | 975.4166667 |

| | | | | |
|---|---|---|---|---|
| sD-MO | 864.0833 | 1062.5 | 900.2143 | 992.0833333 |
| PVL | 972.3333 | 1227.643 | 981.5 | 1075.583333 |
| PreT | 977.25 | 1293.714 | 1045.143 | 1146.083333 |
| preTh_v | 715.0833 | 1062.286 | 861.8571 | 964 |
| vHb | 857.1667 | 1209.5 | 981.3571 | 1084.666667 |
| PreT_a | 993.3333 | 1322.571 | 1050.786 | 1170.25 |
| EmT_r | 847.8333 | 1210.857 | 1001.571 | 1121.083333 |
| Hb | 851.75 | 1186.786 | 939.0714 | 1068.5 |
| dTel | 964.4167 | 1321.929 | 1031.571 | 1162.583333 |
| dTh | 878.1667 | 1211.429 | 986 | 1117.25 |
| Pro | 631.6667 | 898.1429 | 693.1429 | 830.25 |
| dHb | 841.25 | 1149.714 | 887.6429 | 1040.166667 |
| PoR | 573.5 | 832.3571 | 671.6429 | 826.3333333 |
| sPro | 598 | 852 | 641.7143 | 802.1666667 |
| TL | 738.8333 | 1136.75 | 746.9286 | 907.4166667 |
| Tel | 890 | 1217.571 | 930.7143 | 1096.416667 |
| vTel | 758.25 | 1040.571 | 821.7143 | 994.8333333 |
| Ret | 553.6667 | 833.9286 | 580.3571 | 759.5833333 |
| EmT | 879.5 | 1300.714 | 949.6429 | 1153 |
| vTh_a | 752 | 1144.571 | 890.2143 | 1096.333333 |
| OE | 646.8333 | 935 | 678.7857 | 887.75 |
| AF5 | 838.875 | 1163.929 | 961.5 | 1182.541667 |
| vENT | 892.4167 | 1345.714 | 932.0714 | 1168.75 |
| OB | 928.25 | 1241.286 | 851.0714 | 1106.5 |
| Ep | 940.1667 | 1614.464 | 1039.714 | 1478.916667 |
